# Predicting recombination frequency from map distance

**DOI:** 10.1101/2020.12.14.422614

**Authors:** Mikko Kivikoski, Pasi Rastas, Ari Löytynoja, Juha Merilä

## Abstract

Map distance is one of the key measures in genetics and indicates the expected number of crossovers between two loci. Map distance is estimated from the observed recombination frequency using mapping functions, the most widely used of those, Haldane and Kosambi, being developed at the time when the number of markers was low and unobserved crossovers had a substantial effect on the recombination fractions. In contemporary high-density marker data, the probability of multiple crossovers between adjacent loci is negligible and different mapping functions yield the same result, that is, the recombination frequency between adjacent loci is equal to the map distance in Morgans. However, high-density linkage maps contain an interpretation problem: the map distance over a long interval is additive and its association with recombination frequency is not defined. Here, we demonstrate with high-density linkage maps from humans and stickleback fishes that the inverse of Haldane or Kosambi mapping functions fail to predict the recombination frequency from map distance, and show that this is because the expected number of crossovers is not sufficient to predict recombination frequency. We formulate a piecewise function to calculate the probability of no crossovers between the markers that yields more accurate predictions of recombination frequency from map distance. Our results demonstrate that the association between map distance and recombination frequency is context-dependent and no universal solution exists. We anticipate that our study will motivate further research on this subject to yield a more accurate mathematical description of map distance in the context of modern data.

## Introduction

Crossovers in meiosis break the physical linkage among loci and allow formation of recombinant chromosomes and ensure chromosome segregation in meiosis I (Koehler et al. 1996, Hassold et al. 2021). Although crossovers and the resultant recombinations have been studied for more than a hundred years (Sturtevant 1913), many related questions remain unanswered. Due to their profound importance in sexual reproduction, substantial research efforts have focused on better understanding the among-organism and individual variation of crossover rate (e.g., Stapley et al. 2017, Haenel et al. 2018), and on the other hand, recent technologies have been utilized for detecting recombinations at the gamete level (Dréau 2019, Bell et al. 2020, Yang et al. 2022). Crossovers also have implications for statistical properties essential in population genetics, such as the variance of genetic relatedness (Veller et al. 2020). However, one aspect that has gained little attention in the era of high-throughput sequencing is the interpretation of genetic map distances.

Recombinant gametes or offspring can be utilized to build linkage maps that quantify the physical order and map distance (i.e., expected number of crossovers) between loci. Map distances are estimated with mapping functions that attempt to account for the non-additivity of the recombination frequencies due to multiple crossovers between adjacent loci. The two most widely used and referred to mapping functions are probably those of Haldane (1919) and Kosambi (1944, e.g. Lynch and Walsh 1998, Hill and Weir 2011, Otto and Payseur 2019). However, modern sequencing methods and increasing marker density have reduced the utility of these functions in linkage map reconstruction; the probability of multiple crossovers between closely positioned adjacent loci is negligible and all mapping functions yield essentially the same result, *r* = *d*, where *r* is recombination frequency and *d* is map distance in Morgans (e.g., Purcell et al. 2007). As the map distances are estimated only for adjacent loci, they are additive over longer intervals. Consequently, recombination frequency over intermediate or long map distances (e.g., 50 cM) does not follow any simple association.

Although the importance of mapping functions in linkage map construction has decreased over time with increasing access to dense marker data, inverse mapping functions have recently been utilized to predict genetic shuffling in meiosis (Veller et al. 2019) and from that the variance in genetic relatedness (Veller et al. 2020). It is technically trivial to translate map distance to recombination frequency with an inverse of mapping function, but as all mapping functions effectively yield the same map for high-density data, it is not clear which inverse mapping function, if any, would yield the true recombination frequencies. Veller et al. (2020) observed that empirical variance in genetic relatedness among human (*Homo sapiens*) subjects did not match those predicted with the inverse of Kosambi function, indicating that the inverse of Kosambi function is invalid for translating map distances into recombination frequencies.

Here, we show with empirical data from humans and from nine-spined (*Pungitius pungitius*) and three-spined (*Gasterosteus aculeatus*) sticklebacks, that the inverse of Kosambi, Haldane, or linear mapping functions do not translate additive map distances correctly to recombination frequencies. To that end, we propose a new approach to translate map distances to recombination frequencies using a piecewise function based on the probability of no crossovers between the markers. We motivate this approach with empirical data from the nine-spined stickleback and demonstrate its performance with empirical data from humans and nine- and three-spined sticklebacks.

## Materials and methods

### Number of crossovers and recombination frequency

An odd number of gametic crossovers between two loci cause recombination, and the recombination frequency of two loci is equal to the probability of an odd number of gametic crossovers between them. Assuming that there is no chromatid interference, any positive number of crossovers in the bivalent leads to equal proportions of recombinant and non-recombinant gametes, while the absence of crossovers between two loci always leads to non-recombinant gametes. From this follows that recombination frequency *r*, can be expressed as a function of probability of no crossovers between two loci in the bivalent *p*_0_, so that *r* = ½(1-*p*_0_) (Mather 1938, Weeks et al. 2009). To translate map distance into recombination frequency, the association between map distance and *p*_0_ is necessary. Map distance of two loci is the expected number of gametic crossovers between them. However, the distance does not tell the variation around the expectation or the likelihood for no bivalent crossovers (*p*_0_) in that particular interval. Depending on the number of crossovers in the bivalent and their localization, the same map distance can be associated with different values of *p*_0_ and recombination frequency (Fig. 1). To address this ambiguity, we formulate a piecewise function *p*_0_(*k*) that gives the probability of no crossovers between the two loci in the bivalent with *k* crossovers.

**Fig. 1.**
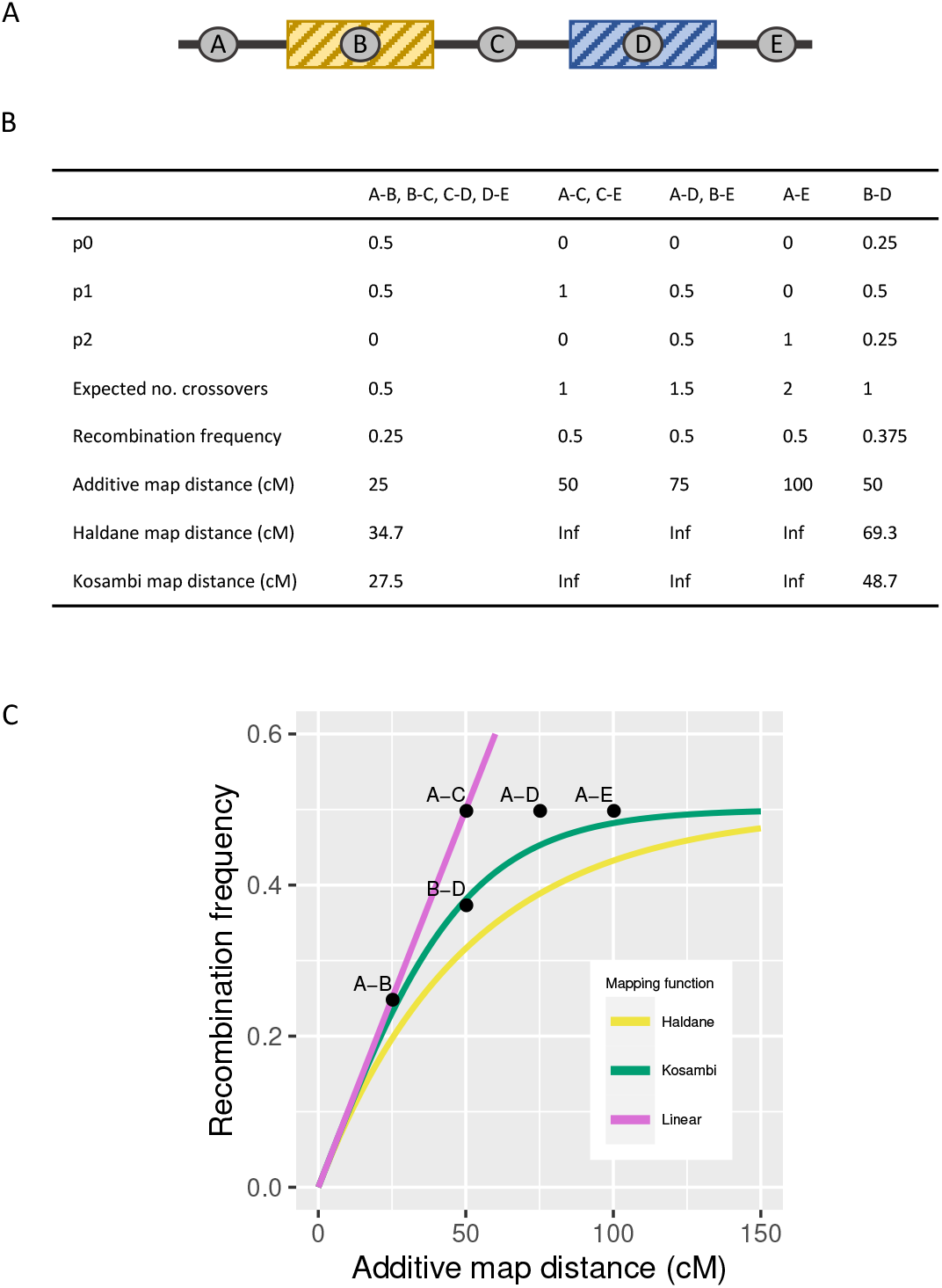
Number and location of crossovers affect the recombination frequency. (A) An example of a chromosome with always two crossovers *in the bivalent* so that one occurs in the yellow and the other one in the blue area. The locations of the crossovers within their distinct regions are independent. (B) The table shows the probability for 0, 1, or 2 crossovers between the markers (*p*_0_, *p*_1_, and *p*_2_, respectively) and the expected number of crossovers *in the bivalent*, the resulting recombination frequency of the marker pair (*in the gamete*) and the estimated map distance for every marker pair using different mapping functions. (C) Graph showing the relationship between the recombination frequency and map distance with different mapping functions. Notably, intervals A-C, A-D, and A-E have the same recombination frequencies but different map distances, whereas marker pairs A-C and B-D have the same map distances but different recombination frequencies.

### Derivation of p_0_(k)

The impetus for the model comes from chromosome 8 (LG8) of the nine-spined stickleback (*Pungitius pungitius*). LG8 is a metacentric chromosome (Kivikoski et al. 2021), where maternal map lengths of the two chromosome arms are 52 and 60 cM, corresponding on average to 1.04 and 1.20 crossovers per bivalent, respectively (Table S1). Out of 934 gametes, 265 have two maternal crossovers and 94% of those gametes have one crossover in both chromosome arms. Because these are gametic crossovers, the number of crossovers in the bivalent could have been two or more. However, the inferred likelihoods for two, three, or four crossovers in the bivalent are 0.85, 0.12, and 0.03, respectively, and likelihoods for other crossover counts are zero (see the section ‘Inference of bivalent crossover rate’ below). Assuming no chromatid interference, the proportion of the gametes with two crossovers descending from meiosis with the same number of crossovers is 0.79 (see supplementary methods). Given this high proportion, those crossovers are, for the sake of example, interpreted to largely represent bivalent crossovers in meiosis with two crossovers. The map positions of the two crossovers are uncorrelated (*r*=0.029, *p*=0.64) and they are distributed approximately uniformly along the map distance (Fig. 2).

**Fig. 2.**
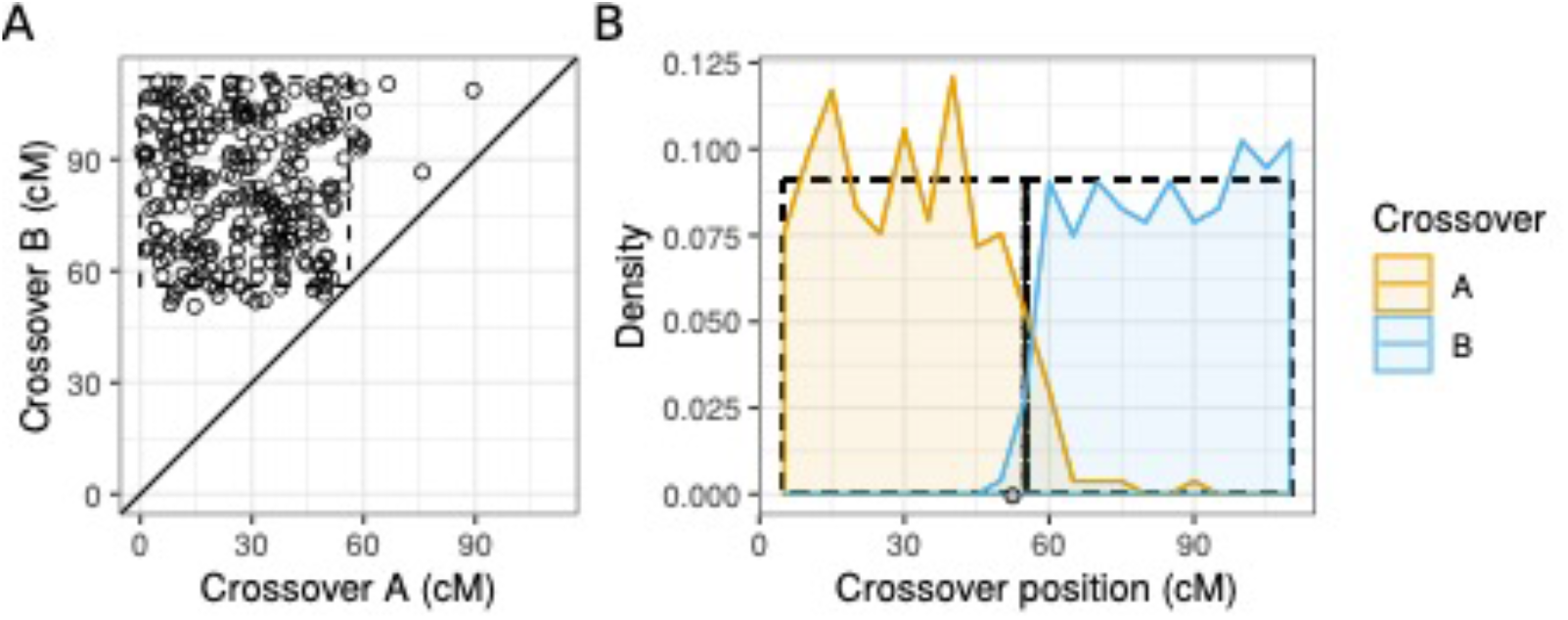
Observed spatial distribution of maternal crossovers in gametes with two maternal crossovers. (A) Each dot corresponds to one offspring (n=265) and coordinates show the map positions of the two crossovers, ‘A’ referring to the crossover closer to chromosome start and ‘B’ to the one closer to chromosome end (*r*=0.029, *p*=0.64). (B) Density plots show the distribution of the two crossovers. Densities are calculated in 5 cM windows. In both panels, the dashed-line rectangles show the expected distribution of crossovers expected under the model presented here.

Putting this observation in a general context, we assume that when there are *k* crossovers in the bivalent they occur in *k* distinct regions, and within those regions the exact localization is independent of other crossovers in the bivalent. Let *d* be the map length of the whole chromosome. If there are *k*(*k* > 0) crossovers in the bivalent, they are assumed to occur in *k* non-overlapping regions of equal size, 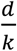, such that every crossover occurs in a different chromosomal region. Within each region, their localizations are uniformly distributed and independent across the regions. For a given marker pair, 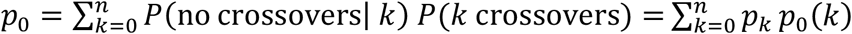, where *n* is the highest possible number of crossovers in the bivalent, *p*_*k*_ is the probability of *k* crossovers in the bivalent and *p*_0_(*k*) is the probability of no crossovers between the markers when there are *k* crossovers in the bivalent. For markers at map positions *m*_*i*_ and *m*_*j*_ (*m*_*j*_ ≥ *m*_*i*_), *p*_0_(*k*) is defined as:

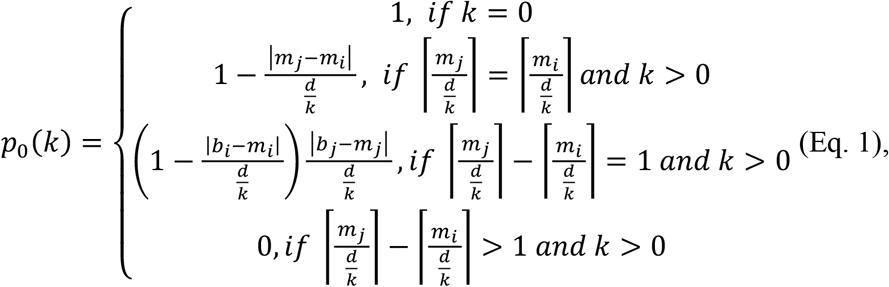

where 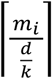 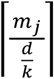 are the crossover regions of the markers *m*_*i*_ and *m*_*j*_, respectively, and *b*_*i*_ and *b*_*j*_ are the upper boundaries for these regions, respectively, so that 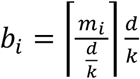 and 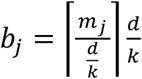. Notation ⌈ ⌉ refers to ceiling function, and 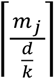 is the least integer greater than, or equal to 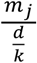.

This function is also applicable for multiple loci. In the case of three markers (*m*_*i*_, *m*_*j*_, *m*_*k*_), the recombination frequencies derived with this function meet the criteria *r*_*ij*_ + *r*_*jk*_ ≥ *r*_*ik*_, where *r*_*ij*_, *r*_*jk*_, and *r*_*ik*_ are the recombination frequencies between markers *m*_*i*_ and *m*_*j*_, *m*_*j*_ and *m*_*k*_, and *m*_*i*_ and *m*_*k*_, respectively (Karlin and Liberman 1978, Weeks 1994; see also Supplementary methods).

Our approach is built on three premises: (1) Absence of chromatid interference; (2) even spacing of crossovers due to crossover interference; and (3) the fact that the number of bivalent crossovers in meiosis I varies across individuals, chromosomes, sexes, and species. The second and the third premises have been demonstrated several times in the literature and are merely facts. Crossover interference causes even distribution of crossovers by their physical distances (micrometers) (Zhang et al. 2014, Zickler and Kleckner 2015) and approximately by base pairs, as the chromatin packing ratio per nucleus is roughly constant (see Fig. S2 in Veller et al. 2019).

However, the spatial distribution per map distance cannot be concluded directly. The number of crossovers per chromosome is not constant but varies between chromosomes, nuclei, individuals, sexes, and species (e.g., Stapley et al. 2017) and hence affects the spatial distribution of crossovers (Charles 1938, Zhang et al. 2014). While the absence of chromatid interference is a common assumption in the literature (e.g., Weinstein 1936, Zhao and Speed 1996, Sandor et al. 2012), this has been empirically tested and confirmed only in a few organisms, including humans and yeast (Zhao et al. 1995, Mancera et al. 2008, Hou et al. 2014, Wang et al. 2019).

In contrast to existing mapping functions, the proposed model is not based on the renewal process (Zhao and Speed 1996) and it does not model crossover interference parametrically. Instead, crossover interference is implemented structurally by assuming that crossovers occur in close proximity less often than would be expected by chance in the absence of crossover interference.

We applied the function for sex-specific recombination data in the 21 chromosomes of nine- and three-spined sticklebacks and in the 22 human autosomes. The total map lengths were derived from the linkage maps and the likelihoods for different numbers of crossovers in the bivalent were inferred from the observed number of crossovers (see ‘Inference of bivalent crossover frequency’ below). Implementation of the *p*_0_(*k*) function in R is provided in the ‘Data archiving’ section.

### Stickleback linkage maps

The crossover frequencies and locations were estimated from the linkage maps described in Kivikoski et al. (2021). For the nine-spined stickleback, high-density linkage maps (22,468 markers informative to conclude crossover) were reconstructed with Lep-MAP3 software (Rastas 2017) from a data set of 133 parents and 938 F_1_ offspring. The parental fish, 46 females and 87 males, were wild-caught individuals from the Baltic Sea coast of Finland (Helsinki, 60°13’N, 25°11’E) that were artificially crossed in laboratory to produce the aforementioned F_1_ offspring (Kivikoski et al. 2021, see also Rastas et al. 2016). Five females were each crossed with a different male, forming five full-sib families, and the other 41 females were each crossed with two different males, which formed 41 half-sib families.

Identification of the single nucleotide polymorphisms (SNPs) of the parental and the F_1_ fish were based on whole-genome sequencing of the parents (5– 10X coverage; Illumina Hiseq platforms, BGI Hong Kong) and DarTseq (Diversity Arrays Technology, Pty Ltd) genotyping of the F_1_ fish (Kivikoski et al. 2021). The read mapping was conducted with BWA-mem (ver. 0.7.15, Li 2013) and the variants were called with SAMtools mpileup (ver 1.9, Li et al. 2009) following the Lep-MAP3 software pipeline (Rastas 2017). The linkage maps were built with Lep-MAP3, and the number of paternal and maternal crossovers were inferred from the observed changes in the haplotype phase of the F_1_ offspring. Crossovers could not be inferred for four crosses with only single offspring, and the final dataset included 934 offspring in total.

The three-spined stickleback linkage maps were based on previously published sequencing data (Pritchard et al. 2017). In short, ninety wild-caught parental fish (Baltic Sea, Helsinki, Finland, 60°13’N, 25°11’E) were crossed such that the males (n=30) were each crossed with two different females (n=60). This yielded 60 and 30 full-sib and half-sib families, respectively, with 517 F_1_ offspring. Genotyping of the parental and the F_1_ fish were based on genotype-by-sequencing, according to the Restriction-site Associated DNA (RAD) sequencing protocol of Elshire et al. (2011). The crossing, rearing protocols and sequencing data are explained in more detail in Leder et al. (2015) and Pritchard et al. (2017). For this study, the RAD reads from Pritchard et al. (2017) were mapped to three-spined stickleback reference genome (v4, Peichel et al. 2017) and the variants were called following the Lep-MAP3 pipeline as for the nine-spined sticklebacks. This yielded 28,187 informative markers to build linkage maps with Lep-MAP3.

For all linkage maps (maternal and paternal maps of the nine- and three-spined stickleback), the genetic distances between the adjacent markers were calculated from recombination frequency with the Haldane mapping function. The distances are additive for non-adjacent markers. There were, on average, 1,070 and 1,342 markers per chromosome in the nine- and three-spined stickleback maps, respectively. Hence, the inter-marker distances were short: on average 19,571 bp corresponding to 0.054 cM and 0.106 cM in the paternal and maternal maps of the nine-spined stickleback, respectively, and 15,461 bp corresponding to 0.043 cM and 0.075 cM in the paternal and maternal maps of the three-spined stickleback, respectively. As all conventional functions yield very similar results for small recombination frequencies, the choice of the mapping function has a minor impact on the map distances.

### Analysis of human recombination data

To evaluate the general applicability of the new function, we analyzed human data from Halldorsson et al. (2019). This consisted of sex-specific linkage maps (their supplementary Data S1 and S2) and the crossover data (their supplementary Data S4). Crossover locations and counts were obtained from the column ‘medsnp’ of the sex-specific linkage maps for the 41,092 probands with both paternal and maternal crossover information. All crossovers were used irrespective of their status regarding the gene conversions (complex, non-complex, or not assessed; see Halldorsson et al. 2019). Moreover, no probands were discarded based on the total number of crossovers in them; the highest number of crossovers per proband per chromosome was 17 maternal crossovers in chromosome 13. The number of markers in the linkage maps ranged from 17,894 to 90,036 depending on the chromosome. We used R (ver. 4.1.1 R Core Team 2018) with seed value 2021 to sample 1.5% of the markers of every chromosome, which yielded 268–1,351 markers (i.e., 35,778–911,925 marker pairs) per chromosome. For every marker pair, we estimated the sex-specific recombination frequency by calculating the proportion of the studied probands (n=41,092) with an odd number of crossovers between the markers.

### Inference of crossover frequency

Assuming there is no chromatid interference, the probability of observing *k* crossovers in a randomly sampled meiotic product depends on the number of crossovers in the bivalent so that 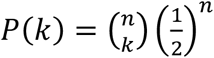, where on is the number of crossovers in the bivalent and 0 ≤ *k* ≤ *n* (Weinstein 1936). The number of crossovers in the bivalent varies not only between the sexes and chromosomes but also between individuals and individual meioses (see Broman and Weber 2000 for an example). Therefore, the observed crossovers in the gamete pool are a sample of crossovers from meioses with a different number of crossovers in the bivalent. As the sampling function above is known, the multinomial distribution of the number of crossovers in the bivalent can be estimated. We used the expectation-maximization (EM) algorithm of Yu and Feingold (2001) to estimate the multinomial distribution for different numbers of crossovers in bivalent. The algorithm approximates the multinomial distribution that maximizes the likelihood of the data consisting of the numbers of meiotic products with 0 … *N* observed crossovers in each chromosome. According to Yu and Feingold (2001), for data where the highest number of observed crossovers in a single meiotic product is *N*, it is sufficient to estimate the multinomial distribution between 0 and 2*N* − 1. The algorithm was applied separately for maternal and paternal crossovers and for each chromosome by pooling all meiotic products (n=934, n=517, n=41,092 for nine- and three-spined stickleback and human data, respectively, Tables S1–S6). The estimated multinomial distribution includes the maximum-likelihood estimate for no crossovers in the bivalent. We applied the bootstrapping test of Yu and Feingold (2001) to estimate if, in case of an estimate above zero, the deviation from that is statistically significant (*p* < 0.05). For the chromosomes with non-significant *p*-values, we used a restricted multinomial distribution that restricts the likelihood of no crossovers to zero for further analyses. This choice was made to assume the obligate crossover as a null hypothesis.

Chromosomes with *p-*value below 0.05 were chr15 of the three-spined stickleback (paternal meioses) and in humans chr21 and chr22 (both maternal and paternal meioses) and chr3 (paternal meioses). For humans, meioses with no crossovers in chromosomes 21 and 22 have been previously reported in cytological studies (e.g. Wang et al. 2017, Hassold et al. 2021), but we are not aware of such findings for the chr3 and further verification is needed. For the sticklebacks, this is the first study testing the obligate crossover hypothesis.

### Performance assessment of functions

The performance of *p*_0_(*k*) and the three inverse mapping functions in predicting recombination frequency from map distance were assessed by calculating the mean absolute error of the predictions and the intra-chromosomal component of genetic shuffling, 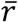 (Veller et al. 2019). The equations for the inverse mapping functions were: 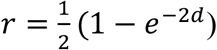 (Haldane 1919), 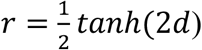 (Kosambi 1944), and 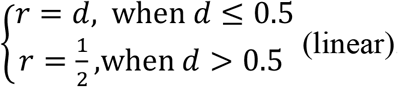. In all functions, *r* is the recombination frequency and *d* is the map distance in Morgans.

The mean absolute error of the predicted recombination frequency was calculated as 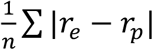, where *r*_*e*_ and *r*_*p*_ are empirical and predicted recombination frequencies per marker pair, respectively, and *n* is the total number of marker pairs. The intra-chromosomal component of 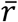 is the part of genetic shuffling due to crossover rate and localization. A higher number of crossovers, and their even distribution increase 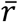, while a low crossover rate and terminal or aggregated localization decrease it. Here, we calculated the intra-chromosomal component of 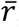, by first predicting the recombination frequency from map distance and then converting it to shuffling according to Eq. 10 of Veller et al. (2019) 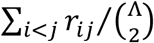, where *r*_*ij*_ is the rate of shuffling, i.e. recombination frequency of locus pair (*i, j*), Λ is the number of loci, and 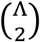 is the number of locus pairs.

## Results

### Map distance and recombination frequency do not have a fixed association

Mapping functions formulate a fixed association between recombination frequency and the expected number of crossovers in a gamete, the map distance. However, the same additive map distance can be associated with different recombination frequencies depending on the crossover positions and the context of the focal loci (Fig. 1). This demonstrates that inverse mapping functions have limitations in predicting recombination frequency from map distance. Maternal crossovers in chromosome 8 of the nine-spined stickleback have a distribution similar to the idealized example (Fig. 2), suggesting that a structural model of crossover localization may be applicable in predicting recombination frequency from map distance.

Another limitation of the mapping functions (and their inverses) is that they do not account for the variation in the number of bivalent crossovers. Inferred probabilities for bivalent crossover counts show that the number of crossovers varies between sexes, among chromosomes, and in meioses in all three studied species (Fig. 3, Tables S1–S6). This further indicates that the inverse of the Kosambi or Haldane mapping function may not be suitable for different species, sexes, or chromosomes.

**Fig. 3.**
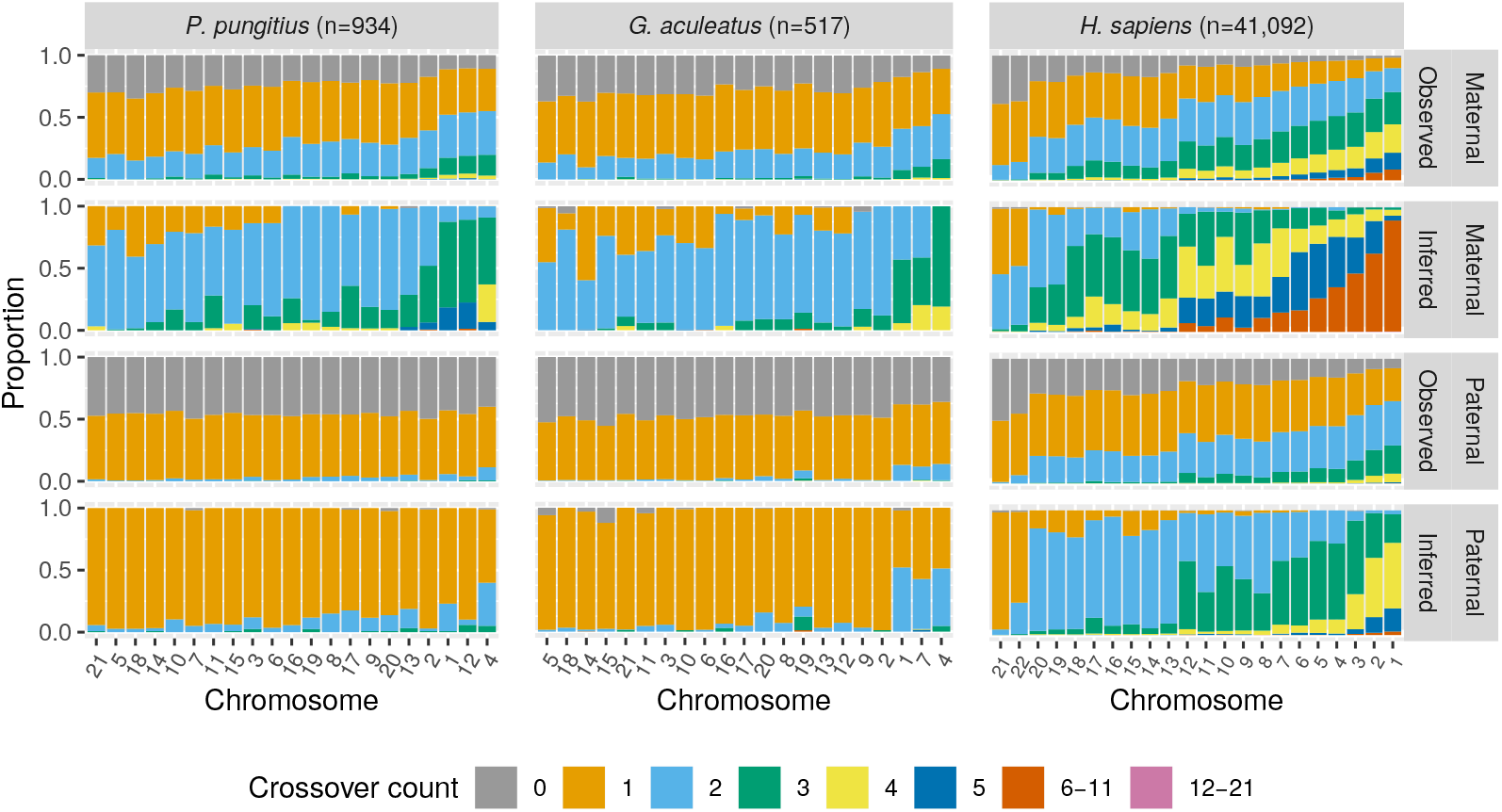
Observed (gametic) and inferred (bivalent) crossover frequency distributions in the nine-spined stickleback (*P. pungitius*), three-spined stickleback (*G. aculeatus*), and human (*H. sapiens*). Each bar shows the proportion of offspring with a certain number of crossovers (Observed) and the inferred proportions of meiosis with a certain number of crossovers in the bivalent in maternal and paternal meioses (Inferred). The inference is based on the expectation-maximization algorithm by Yu and Feingold (2001). Chromosomes are ordered from shortest to longest by length in base pairs. Crossover counts 6–11 and 12–21 are grouped for readability.

### Inverse Kosambi and Haldane mapping functions underestimate recombination frequency

For the three studied species, empirical recombination frequencies lie between the predictions of the inverse Kosambi and linear function (Fig. 4, Fig. S1–S3). This shows that inverse Kosambi and Haldane mapping functions systematically underestimate the recombination frequencies and the linear function overestimates them. The poor performance of the inverse Haldane function is not surprising as it assumes no crossover interference. However, the fact that the inverse Kosambi function, which does implement crossover interference, systematically underestimates recombination frequencies implies that its crossover interference model does not reflect the underlying biology.

**Fig. 4.**
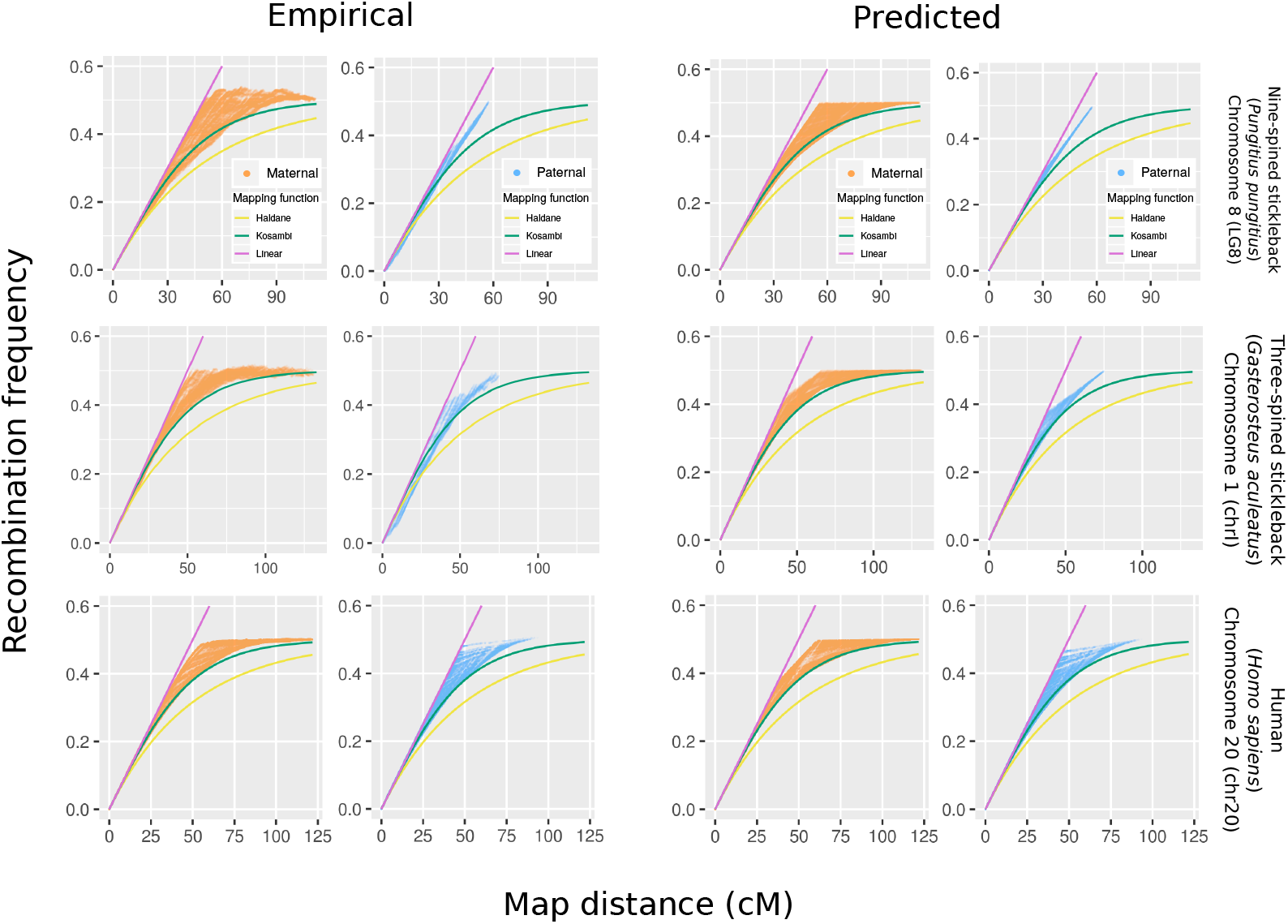
Examples of empirical recombination frequencies (left panels) and predictions by the new function (right) in the nine-spined stickleback (top), three-spined stickleback (middle), and human (bottom). Solid lines show the three inverse mapping functions. Each orange and blue dot is a marker pair in maternal and paternal data, respectively.

### The new p_0_(k) function outperforms the existing functions

We evaluated the new function against existing functions by calculating the mean absolute error of predictions and the intra-chromosomal component of genetic shuffling for the two sexes of three different species. The predictions made with the new *p*_0_(*k*) functions depict the pattern of empirical data (Fig. 4, Fig. S4–S6), and the mean absolute error of those predictions are clearly lowest in five out of the six cases. In nine-spined stickleback males, most meioses have one crossover and the linear function gives a slightly lower error, than *p*_0_(*k*) (Table 1). The excellent overall performance (Table S7–S9) demonstrates that the new function works for different species and on different types of chromosomes. Consistent with the mean absolute error, the new function is superior when assessed on the intra-chromosomal components of 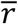 estimated from the empirical and predicted recombination frequencies. For each species and for both sexes, the predictions by the *p*_0_(*k*) function gave results closest to the empirical results (Table 2), demonstrating its potential for the application.

**Table 1.** Means of a per-marker pair absolute error in the predicted recombination frequencies for both sexes of the three study species.

| Species | Sex | Haldane | Kosambi | Linear | $p_0(k)$ |
| --- | --- | --- | --- | --- | --- |
| Human | Paternal | 0.0282 | 0.00936 | 0.00853 | 0.00375 |
|  | Maternal | 0.0257 | 0.00709 | 0.00823 | 0.00346 |
| Nine-spined stickleback | Paternal | 0.0104 | 0.00442 | 0.00267 | 0.00302 |
|  | Maternal | 0.0417 | 0.0152 | 0.0154 | 0.00910 |
| Three-spined stickleback | Paternal | 0.00867 | 0.00399 | 0.00311 | 0.00232 |
|  | Maternal | 0.0438 | 0.0167 | 0.0139 | 0.00981 |

**Table 2.** Intra-chromosomal components of genetic shuffling 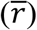 for autosomes. The five columns show 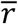 calculated from the empirical recombination frequencies and those predicted by the four functions.

| Species | Sex | Haldane | Kosambi | Linear | $p_0(k)$ | Empirical |
| --- | --- | --- | --- | --- | --- | --- |
| Human | Paternal | 0.012799 | 0.014872 | 0.016796 | 0.015558 | 0.015882 |
|  | Maternal | 0.018091 | 0.020133 | 0.021721 | 0.020672 | 0.020831 |
| Nine-spined stickleback | Paternal | 0.007945 | 0.009261 | 0.01041 | 0.009627 | 0.009819 |
|  | Maternal | 0.01243 | 0.01449 | 0.01646 | 0.01509 | 0.01538 |
| Three-spined stickleback | Paternal | 0.007979 | 0.009292 | 0.010406 | 0.009733 | 0.009648 |
|  | Maternal | 0.011867 | 0.013869 | 0.015800 | 0.014449 | 0.014815 |

Altogether, our analyses show that the inverses of the Kosambi and the Haldane mapping functions systematically underestimate the recombination frequencies. Although the linear function works for chromosomes with an overall crossover rate close to 1 (map length ca. 50 cM), the *p*_0_(*k*) function does not underestimate the recombination frequency to the same extent as the other two functions. Overall, the new *p*_0_(*k*) function gives qualitatively and quantitatively the best results.

## Discussion

### Intrinsic limitations of mapping functions

Map distance tells the expected number of crossovers between two loci and can be used as a proxy of recombination; the longer the map distance the higher the recombination frequency. In principle, translating a map distance to a recombination frequency is trivial, one only needs an inverse of a mapping function such as Kosambi (1944) or Haldane (1919). However, the problem with this approach is that in modern high-density linkage maps, map distances of non-adjacent loci are additive and an inverse mapping function is not guaranteed to give the correct recombination frequency.

The intrinsic limitations of mapping functions, namely that they describe the interference only at a general level and do not account for the variation in the strength of crossover interference or the number of crossovers, have been previously recognized (Crow 1990, Zhao and Speed 1996, Otto and Payseur 2019). However, empirical tests on the performance of inverse mapping functions with modern data are scarce.

### Inverse Kosambi and Haldane mapping functions err with high-density data

Here, we demonstrated that additive map distance can yield an array of different recombination frequencies (Fig. 1) and showed with empirical data from humans and nine- and the three-spined sticklebacks that the inverse of Kosambi and Haldane mapping functions underestimate the recombination frequencies (Fig. 4). We also formulated a new function that outperforms those mapping functions in this task and yields lower error (Table 1) and a more accurate estimate of genetic shuffling (Table 2). The fact that mapping functions fail to predict recombination frequencies implies that those models of crossover interference does not predict crossover localization correctly.

Similar findings were reported by Veller et al. (2020), who showed that the per-chromosome variances of genetic relatedness estimated with the inverse of Kosambi function were higher than those from cytological data that should approximate “true” variance. Our results regarding both sexes of all three studied species were concordant: inverse Kosambi and Haldane functions underestimate recombination frequencies and genetic shuffling 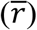, causing overestimation of variance in genetic relatedness, which decreases as a function of 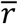 (Veller et al. 2020). Underestimation of recombination frequencies means that the Kosambi map function predicts too many crossovers (excessively long map distance) for observed recombination frequency. The Haldane map function yields even longer map distance, as it assumes no crossover interference.

Stickleback males have primarily one crossover per bivalent (Fig. 3, Table S2, S4) and in those chromosomes the linear function had the best performance (Table S7–S9). The linear function predicts recombination frequency equal to map distance, whereas the *p*_0_(*k*) gives the map distance per the total map length of the chromosome, which yields an underestimate in certain chromosomes. This indicates that the linear function is adequate and should be preferred for chromosomes that mainly have one bivalent crossover, especially in species where this is a norm, such as in Lepidoptera (Davey et al. 2017).

Accurate estimates of recombination frequency are needed in predicting genetic shuffling and from that the variance in genetic relatedness (Visscher et al. 2006). On the other hand, map distance per base pair (cM/Mbp) is used as a measure of recombination in comparative genomics, especially in non-model organisms (Stapley et al. 2017, Martin et al. 2019). This metric is easy to obtain from linkage maps, but as shown here, meaningful interpretation of map distance requires knowledge of the recombination process and interpretation with an inverse mapping function can lead to biased estimates.

### Limitations of the approach

In contrast to existing methods, the approach presented here builds on the fact that the crossover rate varies among organisms and meioses, which leads to differences in spatial distributions of crossover sites, and this variation should be implemented in the model. However, this approach requires knowledge about the probability of different numbers of crossovers in the bivalent, which is not needed by other methods and that cannot be concluded directly from the total map length. These can be inferred from gametic crossovers obtained from linkage maps (as done here) or analysis of haploid offspring (Liu et al. 2015). Alternatively, cytological methods can be used to obtain the bivalent crossover counts directly (Froenicke et al. 2002, Wang et al. 2017). Especially with gametic crossovers that contain the sampling variance caused by the fact that only two chromatids are involved in one crossover event, the sample size must be sufficient to obtain reasonably accurate probability distributions. Implicitly, gametic crossovers were, as a null expectation, assumed to present an unbiased sample of those in the bivalent and that meiotic drive or selection for crossover count has not occurred. However, effects of natural selection could also be involved, for example if the offspring are studied in later life-stages instead of direct investigation of gametes.

In the *p*_0_(*k*) function itself, the most important assumption is the uniform distribution of crossovers per map distance and the arbitrarily defined breakpoints of the “segments”. These assumptions are simplifications that allow mathematically tractable formula. The distribution of crossovers per map distance can be studied from the gametic crossovers only to some extent because they present a subset of those in the bivalent. However, the likelihood for gametes with a certain number of crossovers that show all bivalent crossovers can be calculated and the inherent uncertainty can be estimated (Supplementary methods). Based on the distribution of gametic crossovers per map distance, the assumptions of the model approximate the data in many chromosomes (Fig. S7–S9). However, the non-random distribution of error per map distance shows that all assessed functions have a systematic bias (Fig. S10–S12).

The motivation for the approach presented here came from wild individuals of the nine-spined stickleback, which is not a canonical organism to study crossovers. In contrast to humans for example, the number of crossovers in meiosis varies very little, especially in males. Despite the overall differences in crossover rates (map lengths) of human and sticklebacks, the presented method was demonstrated to work for human data as well.

## Conclusions

The most salient finding of this study is that the inverse of the Kosambi and the Haldane mapping functions systematically underestimate recombination frequencies. Another caveat of using inverse mapping functions to translate additive map distances to recombination frequencies is that they yield one prediction per map distance, which does not match empirical findings.

These findings demonstrate that (intermediate) map distances must be interpreted with care and context-specifically. We also formulated a piecewise function that allows the association between map distance and recombination frequency to be ambiguous. The fact that this function outperforms existing mapping functions in this task indicates that its implementation of crossover interference is more concordant with the data when compared with functions devised earlier. However, the presented function does not replace mapping functions in building linkage maps; it only replaces their inverses in predicting recombination from map distance.

## Acknowledgements

We thank the Biodata Analytics Unit of the University of Helsinki (in particular Jukka Siren) for advice in statistics. We wish to acknowledge CSC – IT Center for Science, Finland, for access to computational resources. Our research was supported by Academy of Finland (grants: 129662, 134728 and 218343 to JM; 322681 to AL; 343656 to PR), the Doctoral Programme in Wildlife Biology Research (LUOVA; funding to MK), the Alfred Kordelin Foundation (grant 210190 to MK), National Natural Science Foundation of China/Research Grants Council (RGC) Joint Research Scheme 2021/2022 (‘N_HKU763/21’ to JM), and support from the Helsinki Institute for Life Sciences (HiLife; grant to JM).

## Author contribution statement

MK formulated the mathematical function introduced here, executed the data analyses, and led the manuscript writing. PR made the stickleback linkage maps and was involved in the formulation of the mathematical function. AL and JM supervised the study and participated in writing the manuscript.

## Data archiving

The stickleback linkage maps and computer code for replication of the analyses of this study are available in Github https://github.com/mikkokivikoski/recombinationStudies. In the repository, the p0kFunction.r file provides an implementation of *p*_0_(*k*) function in R. To facilitate reproducibility of the analyses, analytical steps following the linkage map reconstruction step are integrated into an Anduril2 workflow (Cervera et al. 2019).

## References

Bell, A. D., Mello, C. J., Nemesh, J., Brumbaugh, S. A., Wysoker, A., & McCarroll, S. A. (2020). Insights into variation in meiosis from 31,228 human sperm genomes. Nature, 583: 259–264.

Broman, K. W., & Weber, J. L. (2000). Characterization of human crossover interference. American Journal of Human Genetics, 66(6): 1911–1926.

Cervera, A., Rantanen, V., Ovaska, K., Laakso, M., Nuñez-Fontarnau, J., Alkodsi, A., Casado, J., Facciotto, C., Häkkinen, A., Louhimo, R., Karinen, S., Zhang, K., Lavikka, K., Lyly, L., Pal Singh, M., & Hautaniemi, S. (2019). Anduril 2: upgraded large-scale data integration framework. Bioinformatics, 35(19): 3815–3817.

Charles, D. R. (1938). The spatial distribution of cross-overs in × - chromosome tetrads of Drosophila melanogaster. Journal of Genetics, 36(1): 103–126.

Crow, J. F. (1990). Mapping functions. Genetics, 125(4): 669–67.

Davey, J. W., Barker, S. L., Rastas, P. M., Pinharanda, A., Martin, S. H., Durbin, R., McMillan, W. O., Merrill, R. M., & Jiggins, C. D. (2017). No evidence for maintenance of a sympatric Heliconius species barrier by chromosomal inversions. Evolution Letters, 1(3), 138–154.

Elshire, R. J., Glaubitz, J. C., Sun, Q., Poland, J. A., Kawamoto, K., Buckler, E. S., & Mitchell, S. E. (2011) A robust, simple genotyping-by-sequencing (GBS) approach for high diversity species. PLoS One 6: e19379.

Froenicke, L., Anderson, L. K., Wienberg, J., & Ashley, T. (2002). Male mouse recombination maps for each autosome identified by chromosome painting. American Journal of Human Genetics, 71(6): 1353–1368.

Haenel, Q., Laurentino, T. G., Roesti, M., & Berner, D. (2018). Meta-analysis of chromosome-scale crossover rate variation in eukaryotes and its significance to evolutionary genomics. Molecular Ecology, 27(11): 2477–2497.

Haldane, J. B. S. (1919). The combination of linkage values and the calculation of distances between the loci of linked factors. Journal of Genetics, 8(4): 299–309.

Halldorsson, B. V., Palsson, G., Stefansson, O. A., Jonsson, H., Hardarson, M. T., Eggertsson, H. P., Gunnarsson, B., Oddsson, A., Halldorsson, G. H., Zink, F., Gudjonsson, S. A., Frigge, M. L., Thorleifsson, G., Sigurdsson, A., Stacey, S. N., Sulem, P., Masson, G., Helgason, A., Gudbjartsson, D. F., Thorsteinsdottir, U., & Stefansson, K. (2019). Characterizing mutagenic effects of recombination through a sequence-level genetic map. Science, 363(6425): eaau1043.

Hassold, T., Maylor-Hagen, H., Wood, A., Gruhn, J., Hoffmann, E., Broman, K. W., & Hunt, P. (2021). Failure to recombine is a common feature of human oogenesis. American Journal of Human Genetics, 108(1): 16–24.

Hill, W., & Weir, B. (2011). Variation in actual relationship as a consequence of Mendelian sampling and linkage. Genetics Research, 93(1), 47–64.

Karlin, S., & Liberman, U. (1978). Classifications and comparisons of multilocus recombination distributions. Proceedings of the National Academy of Sciences U.S.A., 75(12): 6332–6336.

Kivikoski, M., Rastas, P., Löytynoja, A., & Merilä, J. (2021). Automated improvement of stickleback reference genome assemblies with Lep-Anchor software. Molecular Ecology Resources, 21(6): 2166–2176.

Koehler, K. E., Hawley, R. S., Sherman, S., & Hassold, T. (1996). Recombination and nondisjunction in humans and flies. Human Molecular Genetics 5(1): 1495–1504.

Kosambi, D. D. (1944). The estimation of map distances from recombination values. Annals of Eugenics, 12(1): 172–175.

Leder, E. H., McCairns, R. J. S., Leinonen, T., Cano, J. M., Viitaniemi, H. M., Nikinmaa, M., Primmer, C. R., & Merilä, J. (2015) The evolution and adaptive potential of transcriptional variation in sticklebacks – signatures of selection and widespread heritability. Molecular Biology and Evolution 32: 674–689.

Li, H., Handsaker, B., Wysoker, A., Fennell, T., Ruan, J., Homer, N., Marth, G., Abecasis, G., & Durbin, R. (2009). The sequence alignment/map format and SAMtools. Bioinformatics, 25: 2078–2079.

Li, H. (2013). Aligning sequence reads, clone sequences and assembly contigs with BWA-MEM. Preprint at http://arxiv.org/abs/1303.3997. p1–3.

Liu, H., Zhang, X., Huang, J., Chen, J. Q., Tian, D., Hurst, L. D., & Yang, S. (2015). Causes and consequences of crossing-over evidenced via a high-resolution recombinational landscape of the honey bee. Genome Biology, 16: 15.

Lynch, M., & Walsh, B. (1998). Genetics and Analysis of Quantitative Traits. Sinauer Associates, Sunderland, MA, pp 393–398.

Martin, S. H., Davey, J. W., Salazar, C., & Jiggins, C. D. (2019). Recombination rate variation shapes barriers to introgression across butterfly genomes. PLoS Biology, 17(2): e2006288.

Mather, K. (1938). Crossing-over. Biological Reviews, 13(3): 252–292.

Otto, S. P., & Payseur, B. A. (2019) Crossover interference: Shedding light on the evolution of recombination. Annual Reviews of Genetics, 53: 19–44.

Peichel, C. L., Sullivan, S. T., Liachko, I., & White, M. A. (2017) Improvement of the threespine stickleback genome using a Hi-C-based proximity-guided assembly. Journal of Heredity, 108: 693–700.

Pritchard, V L., Viitaniemi, H. M., McCairns, R. J. S., Merilä, J., Nikinmaa, M., Primmer, C. R., & Leder, E. H. (2017). Regulatory architecture of gene expression variation in the threespine stickleback Gasterosteus aculeatus. G3: Genes, Genomes, Genetics, 7(1): 165–178.

Purcell, S., Neale, B., Todd-Brown, K., Thomas, L., Ferreira, M. A. R., Bender, D., Maller, J., Sklar, P., de Bakker, P. I. W., Daly, M., & Sham, P. C. (2007). PLINK: A tool set for whole-genome association and population-based linkage analyses. American Journal of Human Genetics, 81(3): 559–575.

Rastas, P., Calboli, F. C. F., Guo, B., Shikano, T., & Merilä, J. (2016). Construction of ultradense linkage maps with Lep-MAP2: Stickleback F2 recombinant crosses as an example. Genome Biology and Evolution, 8(1): 78–93.

Rastas, P. (2017). Lep-MAP3: robust linkage mapping even for low-coverage whole genome sequencing data. Bioinformatics, 33(23): 3726–3732.

R Core Team (2018) R: A Language and Environment for Statistical Computing (R Foundation for Statistical Computing, Vienna, Austria).

Sandor, C., Li, W., Coppieters, W., Druet, T., Charlier, C., & Georges, M. (2012) Genetic Variants in REC8, RNF212, and PRDM9 influence male recombination in cattle. PLoS Genetics, 8(7): e1002854.

Stapley, J., Feulner, P. G. D., Johnston, S. E., Santure, A. W., & Smadja, C. M. (2017). Variation in recombination frequency and distribution across eukaryotes: patterns and processes. Philosophical Transactions of the Royal Society B 372: 20160455.

Sturtevant, A. H. (1915). The behavior of chromosomes as studied through linkage. Z. Abstam. Vererbung, 13: 234–287.

Veller, C., Kleckner, N., & Nowak, M. A. (2019). A rigorous measure of genome-wide genetic shuffling that takes into account crossover positions and Mendel’s second law. Proceedings of National Academy of Science U.S.A. 116 (5) 1659–1668.

Veller, C., Edelman, N. B., Muralidhar, P., & Nowak, M. A. (2020). Variation in genetic relatedness is determined by the aggregate recombination process. Genetics, 216(4): 985–994.

Veller, C., Wang, S., Zickler, D., Zhang, L. & Kleckner, N. (2022). Limitations of gamete sequencing for crossover analysis. Nature 606: E1–E3.

Visscher, P. M., Medland, S. E., Ferreira, M. A. R., Morley, K. I., Zhu, G., Cornes, B. K., Montgomery, G. W., & Martin, N. G. (2006). Assumption-free estimation of heritability from genome-wide identity-by-descent sharing between full siblings. PLoS Genetics 2(3): e41.

Wang, S., Hassold, T., Hunt, P., White, M. A., Zickler, D., Kleckner, N., & Zhang, L. (2017). Inefficient crossover maturation underlies elevated aneuploidy in human female meiosis. Cell, 168(6): 977–989.

Weeks, D. E. (1994) Invalidity of the Rao map function for three loci. Human Heredity, 44: 178–180.

Weinstein, A. (1936). The theory of multiple-strand crossing over. Genetics, 21(3): 155–199.

Yang, L., Gao, Y. Li, M. Park K.-E., Liu, S., Kang, X., Liu, M., Oswalt, A., Fang, L., Telugu, B. P., Sattler, C. G., Li, C., Cole, J. B., Seroussi, E., Xu, L., Yang, L., Zhou, Y., Li, L., Zhang, H., Rosen, B. D., Van Tassell, C. P., Ma, L., & Liu, G. E. (2022). Genome-wide recombination map construction from single sperm sequencing in cattle. BMC Genomics 23: 181.

Yu, K., & Feingold, E. (2001). Estimating the frequency distribution of crossovers during meiosis from recombination data. Biometrics, 57(2), 427–434.

Zhang, L., Liang, Z., Hutchinson, J., & Kleckner, N. (2014) Crossover patterning by the Beam-Film model: Analysis and implications. PLoS Genetics, 10(1): e1004042.

Zhao, H., Speed, T. P. (1996). On genetic map functions. Genetics, 142(4): 1369–1377.

Zickler, D., & Kleckner, N. (2015). Recombination, pairing, and synapsis of homologs during meiosis. Cold Spring Harbor perspectives in biology, 7(6): a016626.

