## Supplementary material for "Predicting recombination frequency from map distance"

### CONTENTS

Supplementary methods
References
Figures S1–S12
Tables S7–S9

### Supplementary methods

#### *Using gametic crossovers to interpret crossover localization in the bivalent*

Linkage maps can be used to infer crossovers and their localization in gametes. However, gametic crossovers are a subset of those in the bivalent and the number of bivalent crossovers in the meiosis that preceded the gamete is generally unknown. This limits the possibilities to study the relationship between crossover count and localization from gametic data[2]. Despite these limitations, the likelihood for bivalent crossover count preceding the gamete can be calculated. Assuming no chromatid interference, the likelihood for a gamete with $k$ crossovers being from a bivalent with $k$ crossovers is:

$$P(CO_{bivalent}=k|CO_{gamete}=k)=\frac{P(CO_{gamete}=k|CO_{bivalent}=k)}{P(CO_{gamete}=k)}=\frac{\frac{\binom{k}{k}}{2^k}p_k}{\sum_{l=k}^{n}\frac{\binom{l}{k}}{2^l}p_l}=\frac{\frac{1}{2^k}p_k}{\sum_{l=k}^{n}\frac{\binom{l}{k}}{2^l}p_l} \tag{1}$$

In Eq. 1 $n$ is the maximum number of bivalent crossovers, $p_k$ and $p_l$ are the likelihoods for $k$ and $l$ bivalent crossovers, respectively. For example, in chromosome 8 (LG8) of the nine-spined stickleback, the inferred probabilities for 2-4 maternal crossovers are 0.85, 0.12 and 0.03, respectively, and probabilities for other crossover counts are 0 (Table S1). The likelihood for a gamete with two maternal crossovers representing the total number of crossovers in meiosis is therefore:

$$\frac{\frac{1}{2^2}p_2}{\sum_{l=2}^{4}\frac{\binom{l}{2}}{2^l}p_l}=\frac{\frac{1}{4}0.85}{\frac{1}{4}0.85+\frac{3}{8}0.12+\frac{6}{16}0.03}=0.79 \tag{2}$$

Tables S1–S6 list the inferred likelihoods for different bivalent crossover counts and for the mode the likelihood that a gamete with the same number of crossovers is equal to the bivalent crossover count.

#### *Proofs for three markers*

According to[1], recombination frequencies $r_{ij}$, $r_{jk}$ and $r_{ik}$ between three markers $i$, $j$ and $k$ at map positions $m_i$, $m_j$ and $m_k$ ($m_i \leq m_j \leq m_k$) should meet the criterion $r_{ij}+r_{jk} \geq r_{ik}$. Here, we show that this holds for the recombination frequencies predicted with the $p_0(k)$ function formulated in this paper. The three cases are (1) all three markers occur in the same crossover region, (2) two markers (here $m_i$ and $m_j$) are in the same region and the third marker is in the next region and (3) all three markers are in three adjacent regions. If there is a marker-free region either between $m_i$ and $m_j$ or $m_j$ and $m_k$, equation $r_{ij}+r_{jk} \geq r_{ik}$ holds, since $r_{ik}=1/2$ and $r_{ij}+r_{jk} \geq 1/2$ as either $r_{ij}$ or $r_{jk}$ is $1/2$ and the other one must be $\geq 0$.

$$\begin{aligned}
\text{Case 1: } m_i, m_j \text{ and } m_k \text{ are in the same crossover region, i.e. } & \left\lceil \frac{m_k}{\frac{d}{k}} \right\rceil = \left\lceil \frac{m_j}{\frac{d}{k}} \right\rceil = \left\lceil \frac{m_i}{\frac{d}{k}} \right\rceil \\
r_{ij}+r_{jk} \geq r_{ik} & \Leftrightarrow \\
\frac{1}{2}\left(1-p_0(k)_{m_i,m_j}\right)+\frac{1}{2}\left(1-p_0(k)_{m_j,m_k}\right) \geq \frac{1}{2}\left(1-p_0(k)_{m_i,m_k}\right) & \Leftrightarrow \\
\frac{1}{2}\left(1-\left(1-\frac{|m_j-m_i|}{\frac{d}{k}}\right)\right)+\frac{1}{2}\left(1-\left(1-\frac{|m_k-m_j|}{\frac{d}{k}}\right)\right) \geq \frac{1}{2}\left(1-\left(1-\frac{|m_k-m_i|}{\frac{d}{k}}\right)\right) & \Leftrightarrow \\
\frac{|m_j-m_i|}{\frac{d}{k}}+\frac{|m_k-m_j|}{\frac{d}{k}} \geq \frac{|m_k-m_i|}{\frac{d}{k}} & \Leftrightarrow \\
|m_j-m_i|+|m_k-m_j| \geq |m_k-m_i| & \Leftrightarrow \\
|m_k-m_i| \geq |m_k-m_i| &
\end{aligned} \tag{3}$$

$$
\begin{aligned}
\text{Case 2: } \left\lceil \frac{m_k}{\frac{d}{k}} \right\rceil - \left\lceil \frac{m_j}{\frac{d}{k}} \right\rceil = 1; \left\lceil \frac{m_j}{\frac{d}{k}} \right\rceil = \left\lceil \frac{m_i}{\frac{d}{k}} \right\rceil \text{ and } b_j = b_i \\
r_{ij} + r_{jk} \geq r_{ik} \Leftrightarrow \\
\frac{1}{2}\left(1 - p_0(k)_{m_i,m_j}\right) + \frac{1}{2}\left(1 - p_0(k)_{m_j,m_k}\right) \geq \frac{1}{2}\left(1 - p_0(k)_{m_i,m_k}\right) \Leftrightarrow \\
\frac{1}{2}\left[1 - \left(1 - \frac{|m_j - m_i|}{\frac{d}{k}}\right)\right] + \frac{1}{2}\left[1 - \left(1 - \frac{|b_j - m_j|}{\frac{d}{k}}\right) \cdot \frac{b_k - m_k}{d_k}\right] \geq \frac{1}{2}\left[1 - \left(1 - \frac{|b_i - m_i|}{\frac{d}{k}}\right) \cdot \frac{|b_k - m_k|}{\frac{d}{k}}\right] \Leftrightarrow ||b_j = b_i \\
1 - 1 + \frac{|m_j - m_i|}{\frac{d}{k}} + 1 - \frac{|b_k - m_k|}{\frac{d}{k}} + \left(\frac{|b_j - m_j|}{\frac{d}{k}} \cdot \frac{|b_k - m_k|}{\frac{d}{k}}\right) \geq 1 - \frac{b_k - m_k}{\frac{d}{k}} + \left(\frac{|b_j - m_i|}{\frac{d}{k}} \cdot \frac{|b_k - m_k|}{\frac{d}{k}}\right) \Leftrightarrow \\
\frac{|m_j - m_i|}{\frac{d}{k}} - \frac{|b_k - m_k|}{\frac{d}{k}} + \left(\frac{|b_j - m_j|}{\frac{d}{k}} \cdot \frac{|b_k - m_k|}{\frac{d}{k}}\right) \geq - \frac{b_k - m_k}{\frac{d}{k}} + \left(\frac{|b_j - m_i|}{\frac{d}{k}} \cdot \frac{|b_k - m_k|}{\frac{d}{k}}\right) \Leftrightarrow \\
\frac{|m_j - m_i|}{\frac{d}{k}} + \left(\frac{|b_j - m_j|}{\frac{d}{k}} \cdot \frac{|b_k - m_k|}{\frac{d}{k}}\right) \geq \left(\frac{|b_j - m_i|}{\frac{d}{k}} \cdot \frac{|b_k - m_k|}{\frac{d}{k}}\right) \Leftrightarrow \\
\frac{|m_j - m_i|}{\frac{d}{k}} \geq \frac{|b_j - m_i|}{\frac{d}{k}} \cdot \frac{|b_k - m_k|}{\frac{d}{k}} - \frac{|b_j - m_j|}{\frac{d}{k}} \cdot \frac{|b_k - m_k|}{\frac{d}{k}} \Leftrightarrow \\
\frac{|m_j - m_i|}{\frac{d}{k}} \geq \left(\frac{|b_j - m_i|}{\frac{d}{k}} - \frac{|b_j - m_j|}{\frac{d}{k}}\right) \cdot \frac{|b_k - m_k|}{\frac{d}{k}} \Leftrightarrow \\
\frac{|m_j - m_i|}{\frac{d}{k}} \geq \frac{|m_j - m_i|}{\frac{d}{k}} \cdot \frac{|b_k - m_k|}{\frac{d}{k}} \Leftrightarrow \\
1 \geq \frac{|b_k - m_k|}{\frac{d}{k}}
\end{aligned} \tag{4}
$$

$$
\begin{aligned}
\text{Case 3: } \left\lceil \frac{m_k}{\frac{d}{k}} \right\rceil - \left\lceil \frac{m_j}{\frac{d}{k}} \right\rceil = 1; \left\lceil \frac{m_j}{\frac{d}{k}} \right\rceil - \left\lceil \frac{m_i}{\frac{d}{k}} \right\rceil = 1 \\
r_{ij} + r_{jk} \geq r_{ik} \Leftrightarrow \\
\frac{1}{2}\left(1 - p_0(k)_{m_i,m_j}\right) + \frac{1}{2}\left(1 - p_0(k)_{m_j,m_k}\right) \geq \frac{1}{2}\left(1 - p_0(k)_{m_i,m_k}\right) \Leftrightarrow \\
\frac{1}{2}\left[1 - \left(1 - \frac{|b_i - m_i|}{\frac{d}{k}}\right) \cdot \frac{|b_j - m_j|}{\frac{d}{k}}\right] + \frac{1}{2}\left[1 - \left(1 - \frac{|b_j - m_j|}{\frac{d}{k}}\right) \cdot \frac{|b_k - m_k|}{\frac{d}{k}}\right] \geq \frac{1}{2}(1-0) \Leftrightarrow \\
1 - \left(1 - \frac{|b_i - m_i|}{\frac{d}{k}}\right) \cdot \frac{|b_j - m_j|}{\frac{d}{k}} + 1 - \left(1 - \frac{|b_j - m_j|}{\frac{d}{k}}\right) \cdot \frac{|b_k - m_k|}{\frac{d}{k}} \geq 1 \Leftrightarrow \\
(-1) \cdot \left(1 - \frac{|b_i - m_i|}{\frac{d}{k}}\right) \cdot \frac{|b_j - m_j|}{\frac{d}{k}} + (-1) \cdot \left(1 - \frac{|b_j - m_j|}{\frac{d}{k}}\right) \cdot \frac{|b_k - m_k|}{\frac{d}{k}} \geq -1 \Leftrightarrow \\
\left(1 - \frac{|b_i - m_i|}{\frac{d}{k}}\right) \cdot \frac{|b_j - m_j|}{\frac{d}{k}} + \left(1 - \frac{|b_j - m_j|}{\frac{d}{k}}\right) \cdot \frac{|b_k - m_k|}{d_k} \leq 1 \\
\text{The maximum value for the terms } \left(1 - \frac{|b_i - m_i|}{\frac{d}{k}}\right) \text{ and } \frac{|b_k - m_k|}{\frac{d}{k}} \text{ is 1.} \\
\text{Therefore, it is sufficient to investigate the equation by setting those values to 1:} \\
1 \cdot \frac{|b_j - m_j|}{\frac{d}{k}} + \left(1 - \frac{|b_j - m_j|}{\frac{d}{k}}\right) \cdot 1 \leq 1 \Leftrightarrow \\
\frac{|b_j - m_j|}{\frac{d}{k}} + 1 - \frac{|b_j - m_j|}{\frac{d}{k}} \leq 1 \Leftrightarrow \\
1 \leq 1
\end{aligned}
\tag{5}
$$

### Supplementary tables S1-S6

Tables S1–S6 (separate csv files) show the number gametes with certain number of crossovers, total map lengths of the chromsosomes and the maximum-likelihood estimates for bivalent crossover counts estimated with the expected-maximization algorithm[3] (see Methods in the main text).

**Table names**

1. Nine-spined stickleback female
2. Nine-spined stickleback male
3. Three-spined stickleback female
4. Three-spined stickleback male
5. Human female
6. Human male

**Column names in the tables**

- n[*]: number of gametes with [*] crossovers
- MLEp[*]: Maximum-likelihood estimate for probability of [*] crossovers in the bivalent.
- MLEpRestricted[*]: Maximum-likelihood estimate for probability of [*] crossovers in the bivalent, by restricting the likelihood for absence of crossovers to 0.
- BootstrapPvalue: Bootstrap $p$-value for the MLEp0. Low $p$-value indicates that non-zero MLEp0 is true finding and meiosis without crossovers may occur.
- MapLength: Map length (cM) of the chromosome.
- MapLengthParm: Map length (cM) of the shorter chromosome arm.
- MapLengthQarm: Map length (cM) of the longer chromosome arm.
- ModeBivalent: Inferred mode of number of crossovers in the bivalent (see columns MLEpRestricted[*]).
- PgameteMode: Probability that gamete with the 'ModeBivalent' crossover count descends from meiosis with the same number of crossovers (see Eq. 1 in supplementary methods).

### References


**1.** S. Karlin and U. Liberman. Classifications and comparisons of multilocus recombination distributions. *Proceedings of the National Academy of Sciences USA*, 75(12):6332–6336, 1978.

**2.** C. Veller, S. Wang, D. Zickler, L. Zhang, and N. Kleckner. Limitations of gamete sequencing for crossover analysis. *Nature*, 606:E1–E3, 2022.

**3.** K. Yu and E. Feingold. Estimating the frequency distribution of crossovers during meiosis from recombination data. *Biometrics*, 57(2):427–434, 2001.

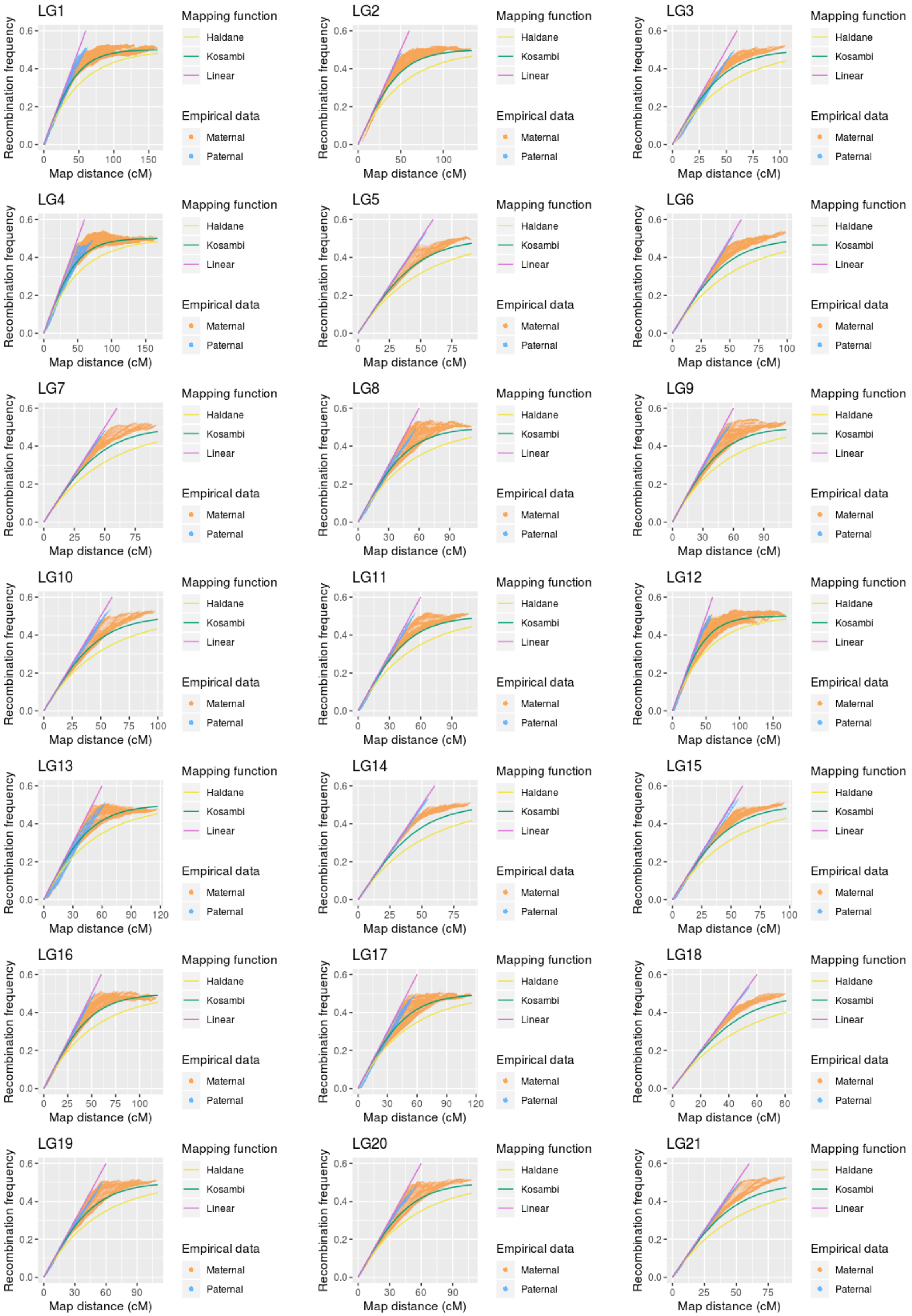


**Figure S1.** Empirical recombination frequencies in the 21 nine-spined stickleback chromosomes. Solid lines show the three inverse mapping functions. Each orange and blue dot is a marker pair in maternal and paternal data, respectively.

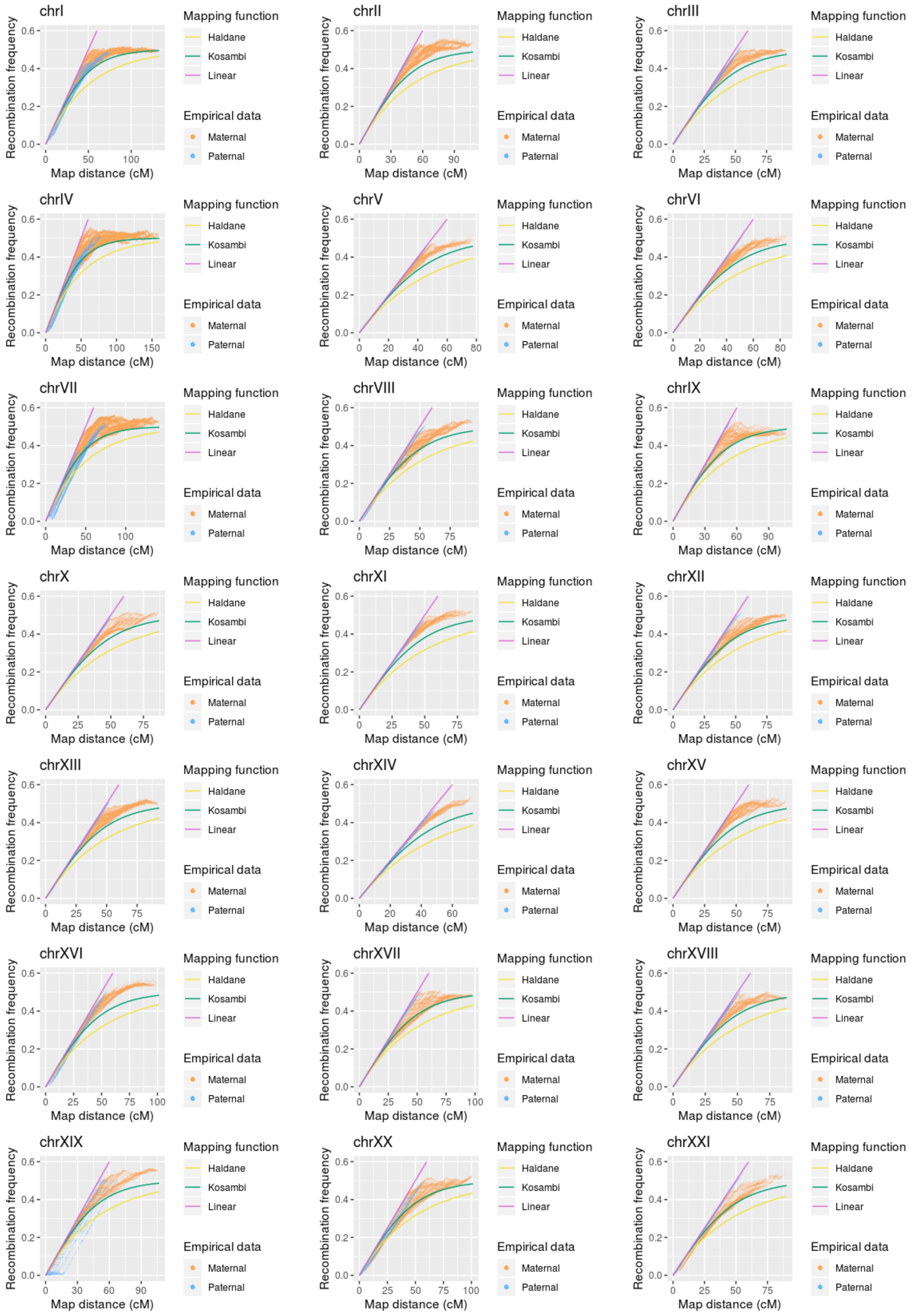


**Figure S2.** Empirical recombination frequencies in the 21 three-spined stickleback chromosomes. Solid lines show the three inverse mapping functions. Each orange and blue dot is a marker pair in maternal and paternal data, respectively.

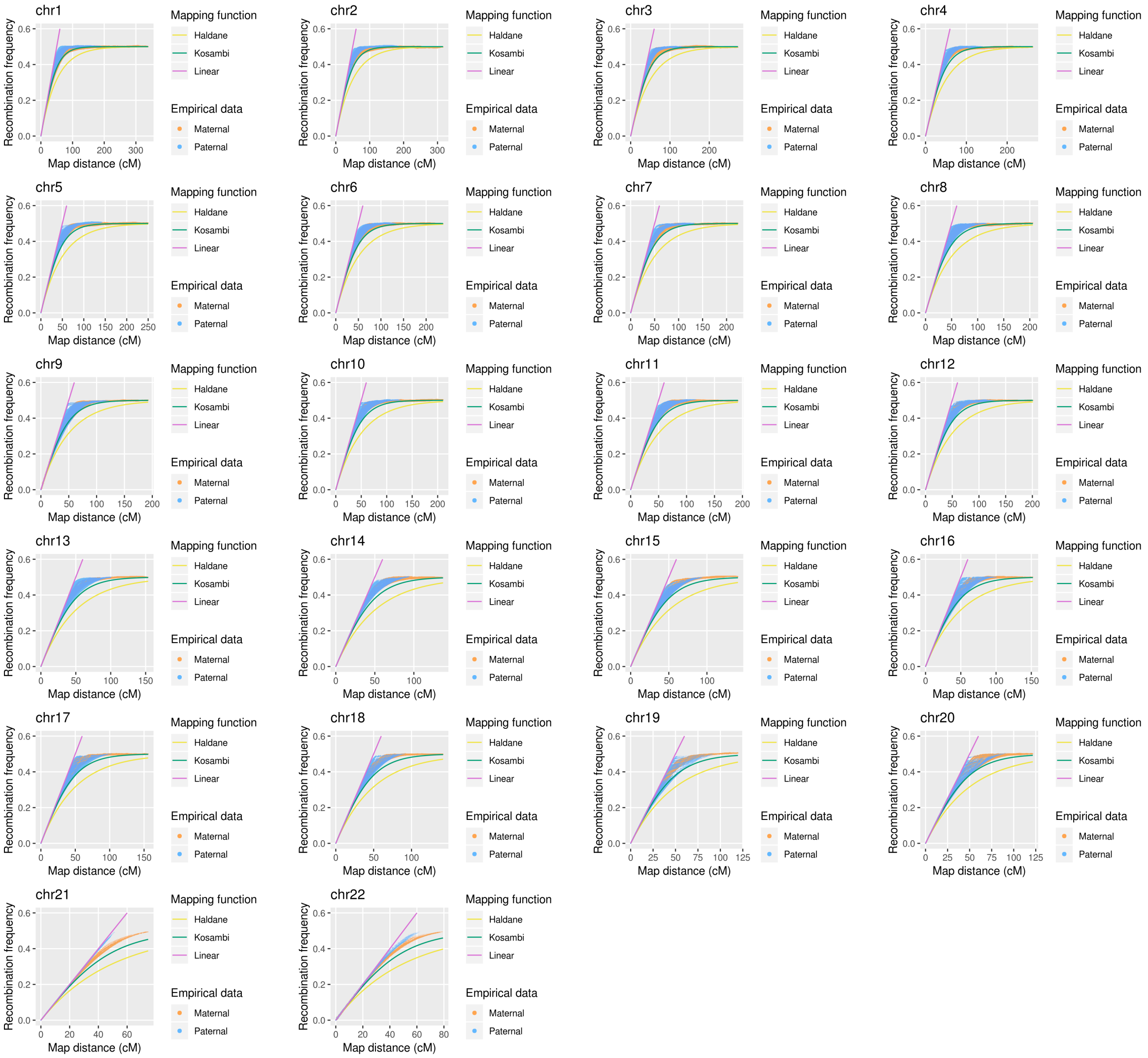


**Figure S3.** Empirical recombination frequencies in the 22 human autosomes. Solid lines show the three inverse mapping functions. Each orange and blue dot is a marker pair in maternal and paternal data, respectively.

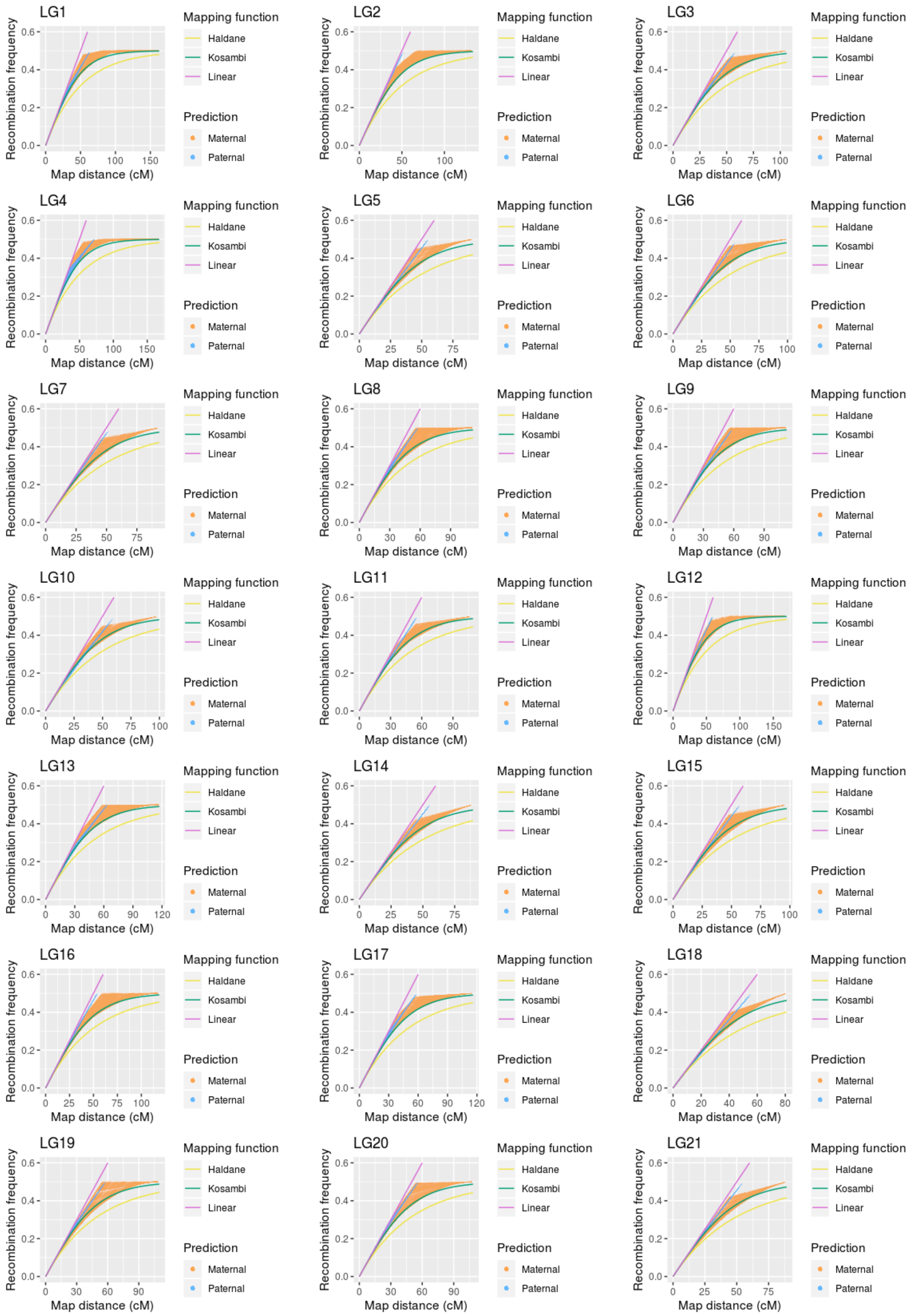


**Figure S4.** Recombination frequencies predicted with the $p_0(k)$ function for the 21 nine-spined stickleback chromosomes.

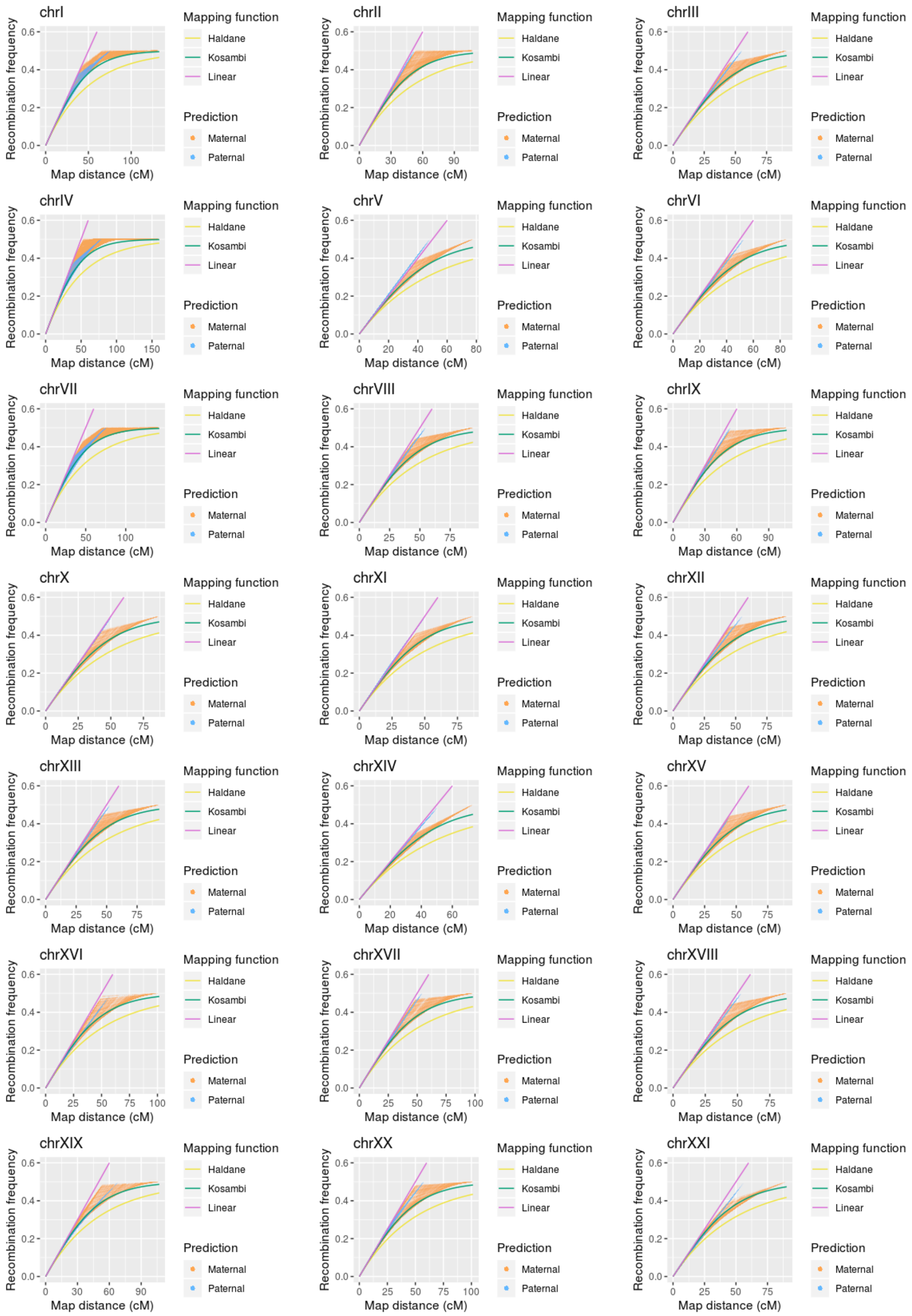


**Figure S5.** Recombination frequencies predicted with the $p_0(k)$ function for the 21 three-spined stickleback chromosomes.

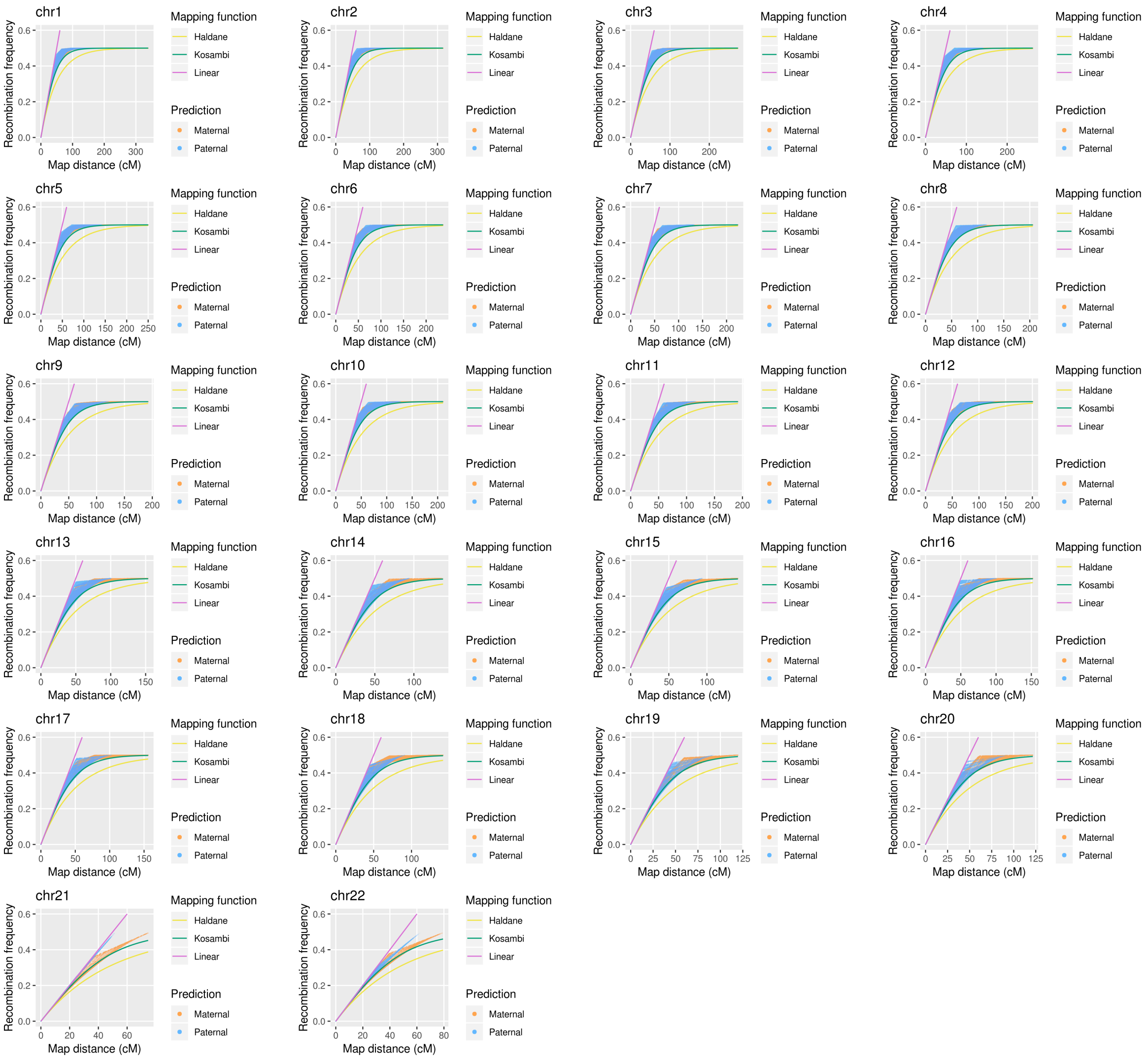


**Figure S6.** Recombination frequencies predicted with the $p_0(k)$ function for the 22 human autosomes.

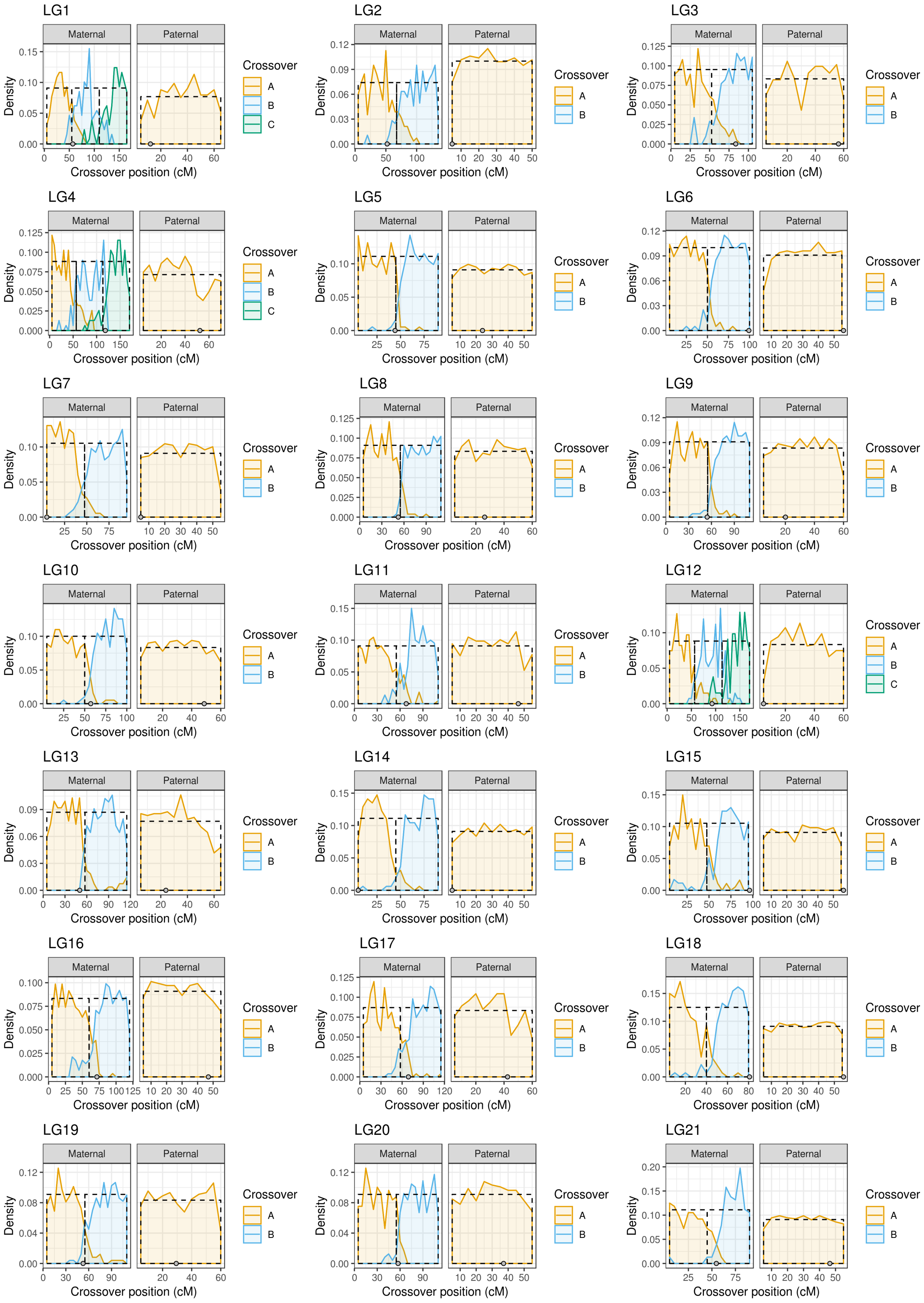


**Figure S7.** Empirical distributions of gametic crossovers per map distance in the 21 nine-spined stickleback chromosomes. Distributions are calculated from gametes whose crossover count is equal to the mode of the bivalent crossover count. Letters A–C refer to the order of the crossovers. Dashed rectangles show the assumed distributions of the crossovers as implemented in the $p_0(k)$ function. Centromeres are shown with the grey dots

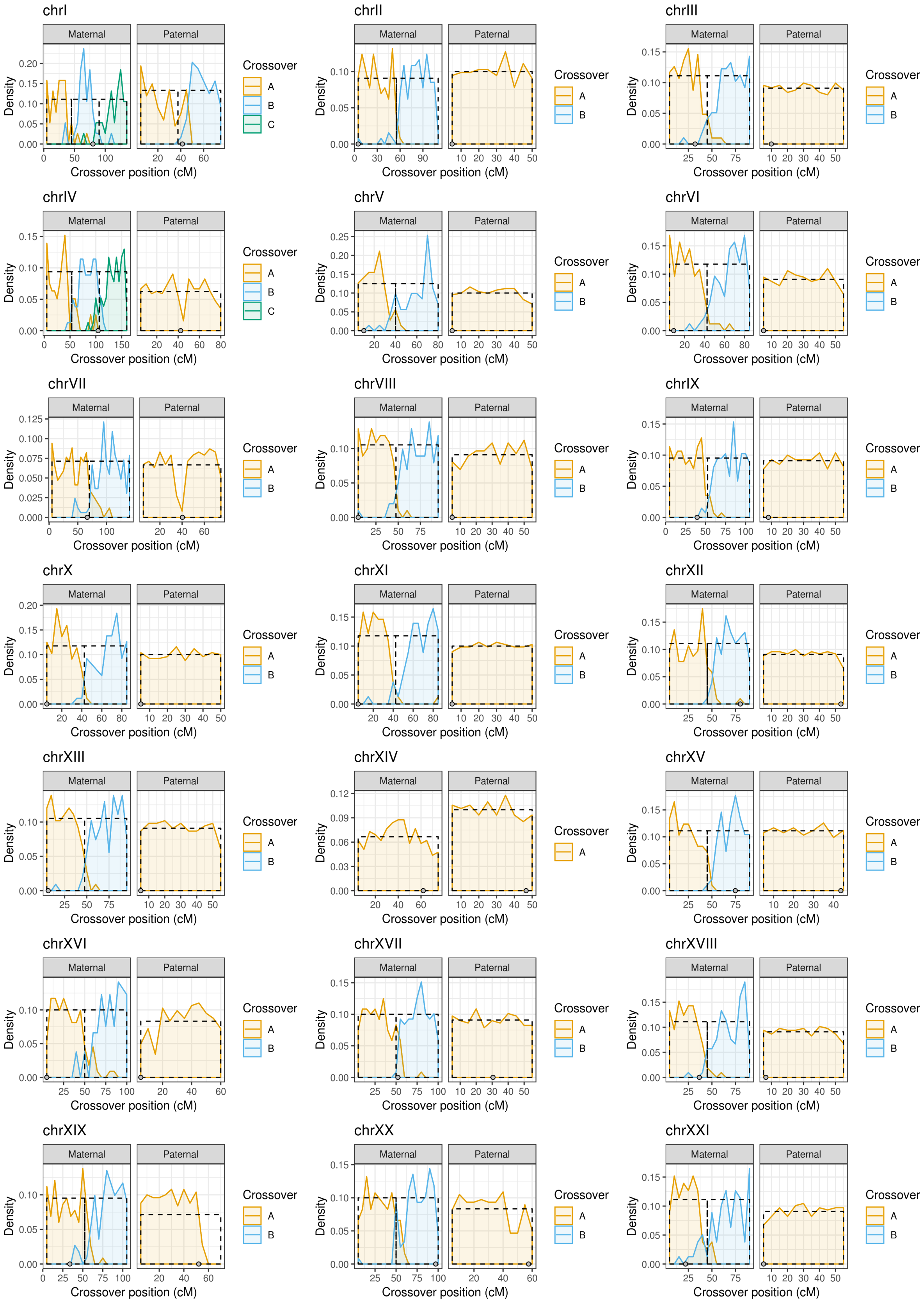


**Figure S8.** Empirical distributions of gametic crossovers per map distance in the 21 three-spined stickleback chromosomes. Distributions are calculated from gametes whose crossover count is equal to the mode of the bivalent crossover count. Letters A–C refer to the order of the crossovers. Dashed rectangles show the assumed distributions of the crossovers as implemented in the $p_0(k)$ function. Centromeres are shown with the grey dots.

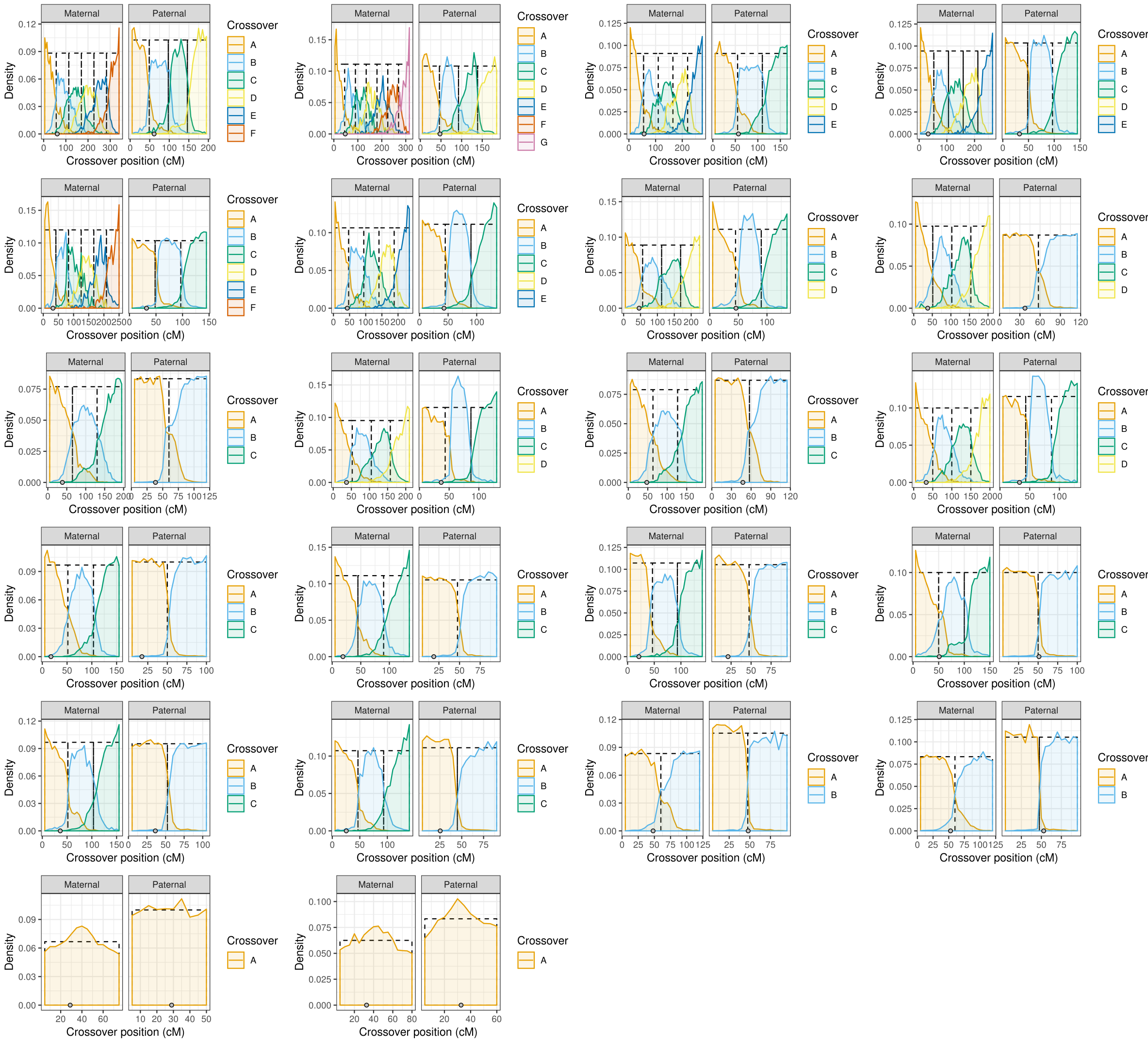


**Figure S9.** Empirical distributions of gametic crossovers per map distance in the 22 human chromosomes. Distributions are calculated from gametes whose crossover count is equal to the mode of the bivalent crossover count. Letters A–G refer to the order of the crossovers. Dashed rectangles show the assumed distributions of the crossovers as implemented in the $p_0(k)$ function. Centromeres are shown with the grey dots.

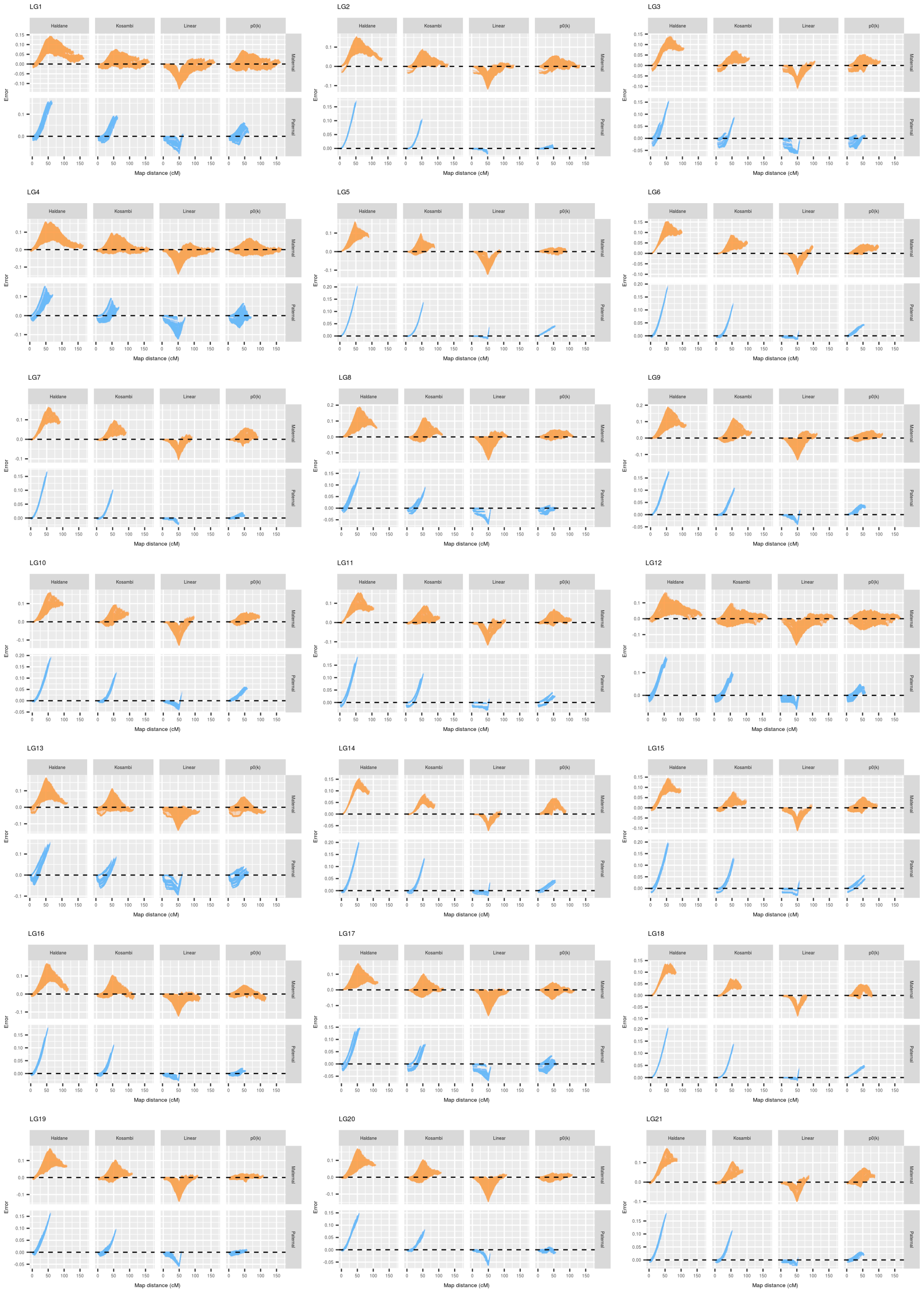


**Figure S10.** Error, i.e. the differences between empirical and predicted recombination frequencies with the different functions for the 21 nine-spined stickleback chromosomes.

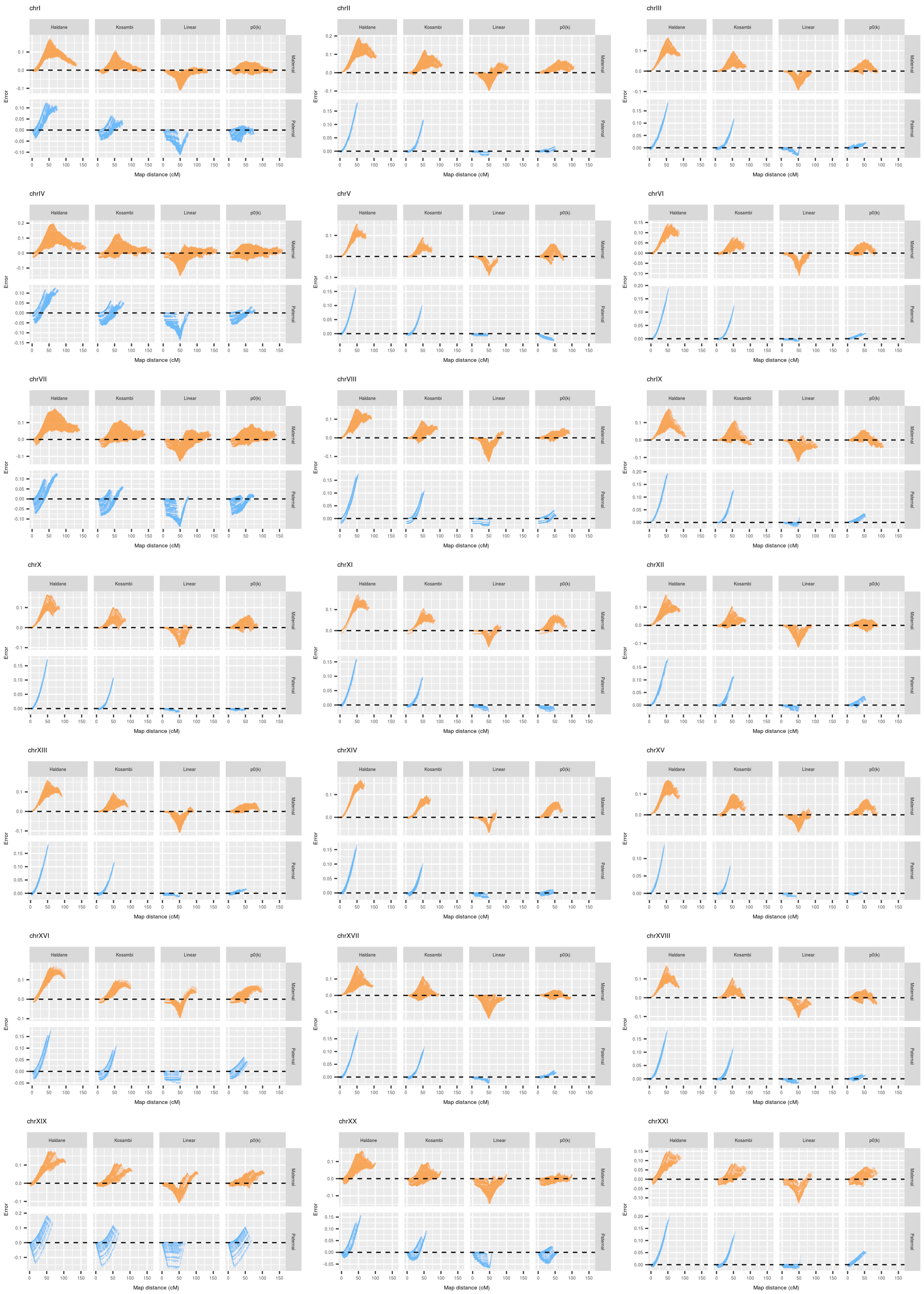


**Figure S11.** Error, i.e. the differences between empirical and predicted recombination frequencies with the different functions for the 21 three-spined stickleback chromosomes.

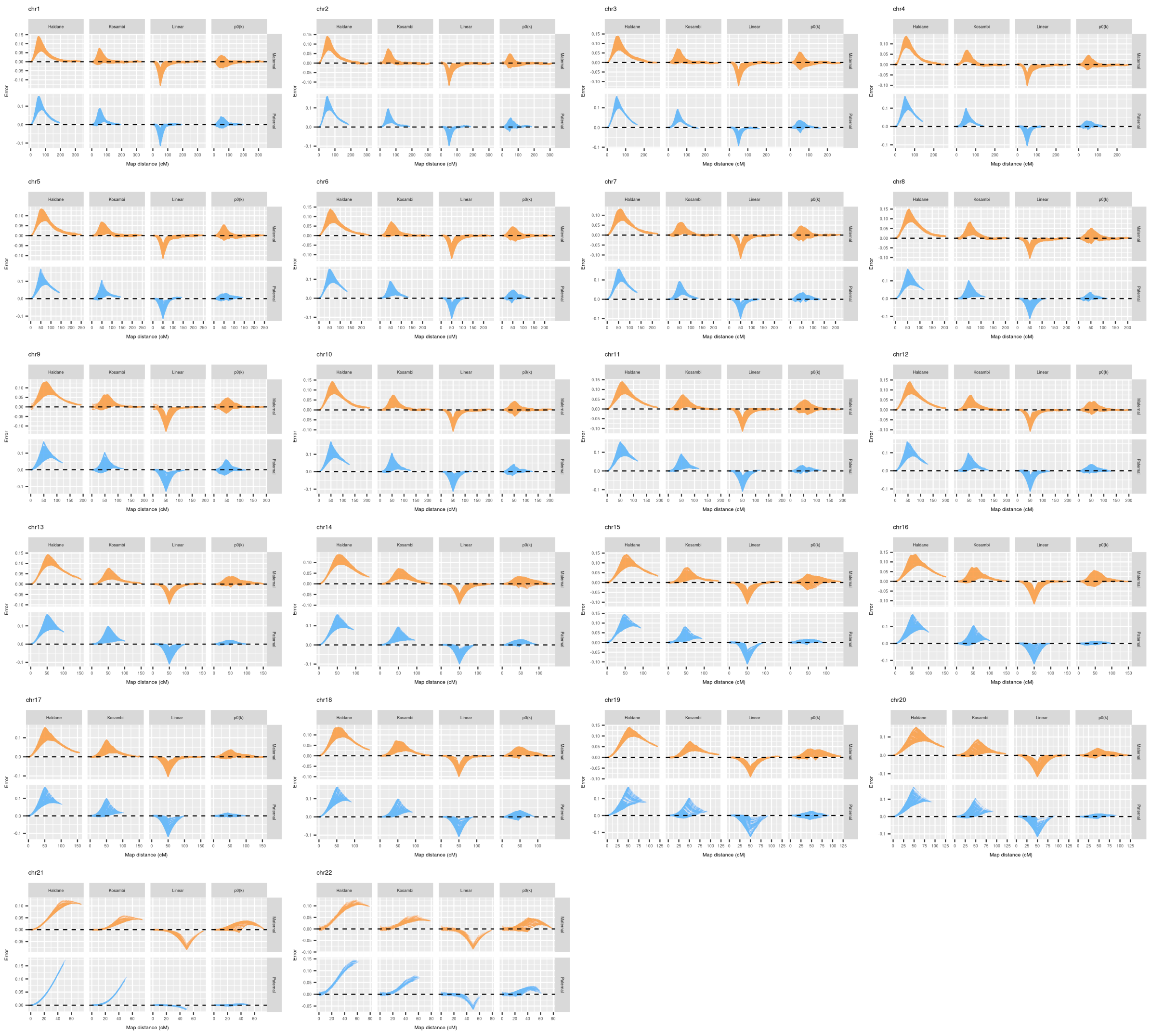


**Figure S12.** Error, i.e. the differences between empirical and predicted recombination frequencies with the different functions for the 22 human chromosomes.

**Table S7.** Mean absolute error of the predicted recombination frequencies in the nine-spined stickleback. Lowest absolute values per sex and chromosome are in bold type.

| | Maternal | | | | Paternal | | | |
|---|---|---|---|---|---|---|---|---|
| Chromosome | Haldane | Kosambi | Linear | $p_0(k)$ | Haldane | Kosambi | Linear | $p_0(k)$ |
| LG1 | 0.0466 | 0.015 | 0.0193 | **0.0095** | 0.0083 | 0.0039 | **0.0023** | 0.0032 |
| LG2 | 0.0478 | 0.0166 | 0.0145 | **0.0103** | 0.0084 | 0.0034 | **0.0004** | 0.0005 |
| LG3 | 0.0378 | 0.0127 | 0.0155 | **0.0096** | 0.0091 | 0.004 | 0.0068 | **0.0028** |
| LG4 | 0.0502 | 0.0183 | 0.0169 | **0.0115** | 0.0069 | 0.0031 | 0.0055 | **0.0030** |
| LG5 | 0.0342 | 0.0115 | 0.013 | **0.0023** | 0.0142 | 0.006 | **0.0006** | 0.0046 |
| LG6 | 0.0439 | 0.0168 | 0.0112 | **0.0086** | 0.0112 | 0.0048 | **0.0007** | 0.0041 |
| LG7 | 0.0406 | 0.0148 | 0.012 | **0.0079** | 0.0111 | 0.0047 | **0.0008** | 0.0013 |
| LG8 | 0.0456 | 0.0188 | 0.0134 | **0.0088** | 0.0079 | 0.0030 | 0.0040 | **0.0018** |
| LG9 | 0.0405 | 0.0163 | 0.0127 | **0.0065** | 0.0139 | 0.0055 | **0.0021** | 0.0042 |
| LG10 | 0.0378 | 0.0137 | 0.0139 | **0.0085** | 0.0124 | 0.0051 | **0.0025** | 0.0052 |
| LG11 | 0.0414 | 0.013 | 0.0156 | **0.0097** | 0.0108 | 0.0048 | **0.0017** | 0.0028 |
| LG12 | 0.043 | 0.0167 | 0.0216 | **0.0142** | 0.0094 | 0.0042 | **0.0021** | 0.0028 |
| LG13 | 0.0387 | 0.0149 | 0.0175 | **0.0102** | 0.0098 | 0.0043 | 0.0062 | **0.0040** |
| LG14 | 0.0394 | 0.0162 | **0.0074** | 0.0119 | 0.0152 | 0.0068 | **0.0008** | 0.0047 |
| LG15 | 0.0363 | 0.0117 | 0.0168 | **0.0085** | 0.0127 | 0.0056 | **0.0012** | 0.0045 |
| LG16 | 0.0424 | 0.0152 | 0.0155 | **0.0085** | 0.0106 | 0.0043 | 0.0017 | **0.0013** |
| LG17 | 0.0376 | 0.0139 | 0.0193 | **0.0085** | 0.0094 | 0.0043 | 0.0040 | **0.0029** |
| LG18 | 0.0346 | 0.0127 | 0.0095 | **0.0072** | 0.0158 | 0.0072 | **0.0003** | 0.0058 |
| LG19 | 0.0384 | 0.0136 | 0.0154 | **0.00400** | 0.0098 | 0.0033 | 0.0039 | **0.0011** |
| LG20 | 0.0378 | 0.015 | 0.0155 | **0.0045** | 0.0097 | 0.0033 | 0.0025 | **0.0012** |
| LG21 | 0.0365 | 0.0136 | 0.0110 | **0.0108** | 0.0126 | 0.0053 | **0.0011** | 0.0026 |

**Table S8.** Mean absolute error of the predicted recombination frequencies in the three-spined stickleback. Lowest absolute values per sex and chromosome are in bold type.

| | Maternal | | | | Paternal | | | |
|---|---|---|---|---|---|---|---|---|
| Chromosome | Haldane | Kosambi | Linear | $p_0(k)$ | Haldane | Kosambi | Linear | $p_0(k)$ |
| chrI | 0.0479 | 0.0161 | 0.015 | **0.0087** | 0.0081 | 0.0033 | 0.0061 | **0.0021** |
| chrII | 0.0541 | 0.0237 | **0.0107** | 0.0128 | 0.0086 | 0.0036 | **0.0005** | 0.0008 |
| chrIII | 0.0394 | 0.0146 | 0.0103 | **0.0075** | 0.0111 | 0.0042 | 0.0023 | **0.0015** |
| chrIV | 0.0533 | 0.0207 | 0.0165 | **0.0108** | 0.0060 | 0.0034 | 0.0063 | **0.0025** |
| chrV | 0.0358 | 0.0131 | 0.0094 | **0.0088** | 0.0095 | 0.0037 | **0.0002** | 0.0024 |
| chrVI | 0.0373 | 0.0127 | 0.0125 | **0.0084** | 0.0086 | 0.0034 | **0.0004** | 0.0015 |
| chrVII | 0.0507 | 0.0209 | 0.0175 | **0.0154** | 0.0077 | 0.0058 | 0.0108 | **0.005** |
| chrVIII | 0.0413 | 0.0141 | 0.0156 | **0.0067** | 0.0092 | 0.0041 | **0.0012** | 0.0017 |
| chrIX | 0.0406 | 0.0142 | 0.0145 | **0.0071** | 0.0109 | 0.0047 | **0.0002** | 0.0024 |
| chrX | 0.0375 | 0.0133 | 0.0113 | **0.008** | 0.010 | 0.0038 | 0.0009 | **0.0007** |
| chrXI | 0.0443 | 0.0189 | **0.0071** | 0.0159 | 0.0082 | 0.0032 | **0.0007** | 0.0014 |
| chrXII | 0.0393 | 0.0121 | 0.0168 | **0.0046** | 0.0073 | 0.0029 | **0.0006** | 0.0014 |
| chrXIII | 0.0444 | 0.0162 | 0.0126 | **0.008** | 0.0085 | 0.0035 | **0.0007** | 0.001 |
| chrXIV | 0.036 | 0.0154 | **0.0055** | 0.013 | 0.0111 | 0.0044 | **0.0008** | 0.0011 |
| chrXV | 0.0479 | 0.0204 | **0.0078** | 0.0127 | 0.0055 | 0.0018 | 0.0006 | **0.0003** |
| chrXVI | 0.0465 | 0.0203 | **0.0119** | 0.013 | 0.0106 | 0.005 | **0.0025** | 0.0047 |
| chrXVII | 0.0371 | 0.0125 | 0.0165 | **0.0042** | 0.0103 | 0.004 | **0.0012** | 0.0014 |
| chrXVIII | 0.0364 | 0.0125 | 0.0113 | **0.0058** | 0.0105 | 0.0042 | 0.0012 | **0.0010** |
| chrXIX | 0.0473 | 0.019 | 0.0181 | **0.0107** | 0.0102 | 0.0069 | 0.00675 | **0.00673** |
| chrXX | 0.0382 | 0.0154 | 0.022 | **0.0066** | 0.0067 | 0.0036 | 0.0041 | **0.0030** |
| chrXXI | 0.0307 | **0.01040** | 0.0185 | 0.01041 | 0.0102 | 0.0043 | **0.0005** | 0.0046 |

**Table S9.** Mean absolute error of the predicted recombination frequencies in human. Lowest absolute values per sex and chromosome are in bold type.

| | Maternal | | | | Paternal | | | |
|---|---|---|---|---|---|---|---|---|
| Chromosome | Haldane | Kosambi | Linear | $p_0(k)$ | Haldane | Kosambi | Linear | $p_0(k)$ |
| chr1 | 0.021 | 0.0056 | 0.007 | **0.003** | 0.0317 | 0.0108 | 0.0072 | **0.0044** |
| chr2 | 0.0213 | 0.0051 | 0.0081 | **0.0028** | 0.0322 | 0.0111 | 0.0072 | **0.004** |
| chr3 | 0.024 | 0.0062 | 0.0082 | **0.0036** | 0.0317 | 0.0108 | 0.0087 | **0.0044** |
| chr4 | 0.024 | 0.006 | 0.0085 | **0.0028** | 0.0305 | 0.0103 | 0.0092 | **0.0039** |
| chr5 | 0.0257 | 0.0068 | 0.008 | **0.0029** | 0.0307 | 0.0107 | 0.0089 | **0.0037** |
| chr6 | 0.0264 | 0.0071 | 0.0082 | **0.0034** | 0.0282 | 0.0092 | 0.0096 | **0.0045** |
| chr7 | 0.0265 | 0.007 | 0.0087 | **0.0035** | 0.0289 | 0.0093 | 0.0099 | **0.0043** |
| chr8 | 0.0269 | 0.0073 | 0.0088 | **0.0037** | 0.0249 | 0.0074 | 0.0101 | **0.003** |
| chr9 | 0.0276 | 0.0076 | 0.0088 | **0.0038** | 0.0251 | 0.0079 | 0.0112 | **0.0056** |
| chr10 | 0.028 | 0.0081 | 0.0078 | **0.0033** | 0.0265 | 0.0084 | 0.0097 | **0.0036** |
| chr11 | 0.0285 | 0.0081 | 0.0087 | **0.0043** | 0.0243 | 0.0071 | 0.0099 | **0.0033** |
| chr12 | 0.0286 | 0.0082 | 0.0084 | **0.0034** | 0.0269 | 0.0088 | 0.0092 | **0.0033** |
| chr13 | 0.031 | 0.0098 | 0.0092 | **0.0043** | 0.0253 | 0.0091 | 0.0071 | **0.0026** |
| chr14 | 0.0298 | 0.009 | 0.0103 | **0.0046** | 0.0262 | 0.0098 | 0.0065 | **0.0036** |
| chr15 | 0.03 | 0.009 | 0.0098 | **0.005** | 0.0223 | 0.0068 | 0.0084 | **0.0021** |
| chr16 | 0.0297 | 0.0093 | 0.0076 | **0.0042** | 0.0236 | 0.0081 | 0.0081 | **0.0014** |
| chr17 | 0.0308 | 0.0099 | 0.0078 | **0.0036** | 0.0212 | 0.0061 | 0.0088 | **0.0016** |
| chr18 | 0.0318 | 0.0106 | 0.0084 | **0.0043** | 0.0204 | 0.0065 | 0.0062 | **0.003** |
| chr19 | 0.0321 | 0.0114 | 0.0078 | **0.0051** | 0.0181 | 0.0059 | 0.007 | **0.0021** |
| chr20 | 0.031 | 0.0103 | 0.0086 | **0.0037** | 0.0197 | 0.0065 | 0.0065 | **0.0013** |
| chr21 | 0.022 | 0.0072 | 0.0074 | **0.0049** | 0.0157 | 0.0062 | 0.0007 | **0.0006** |
| chr22 | 0.0239 | 0.0078 | 0.0086 | **0.0054** | 0.0173 | 0.0064 | **0.0024** | 0.0027 |
